# The genome-wide, multi-layered architecture of chromosome pairing in early *Drosophila* embryos

**DOI:** 10.1101/443028

**Authors:** Jelena Erceg, Jumana AlHaj Abed, Anton Goloborodko, Bryan R. Lajoie, Geoffrey Fudenberg, Nezar Abdennur, Maxim Imakaev, Ruth B. McCole, Son C. Nguyen, Wren Saylor, Eric F. Joyce, T. Niroshini Senaratne, Mohammed A. Hannan, Guy Nir, Job Dekker, Leonid A. Mirny, Chao-ting Wu

## Abstract

Genome organization involves *cis* and *trans* chromosomal interactions, both implicated in gene regulation, development, and disease. Here, we focused on *trans* interactions in *Drosophila*, where homologous chromosomes are paired in somatic cells from embryogenesis through adulthood. We first addressed the long-standing question of whether pairing extends genome-wide and, to this end, developed a haplotype-resolved Hi-C approach that uses a new strategy to minimize homolog misassignment and thus robustly distinguish *trans*-homolog from *cis* contacts. This approach revealed striking genome-wide pairing in *Drosophila* embryos. Moreover, we discovered pairing to be surprisingly structured, with *trans*-homolog domains and interaction peaks, many coinciding with the positions of analogous *cis* features. We also found a significant correlation between pairing and the chromatin accessibility mediated by the pioneer factor Zelda. Our findings reveal a complex, highly structured organization underlying homolog pairing, first discovered more than a century ago.

**One Sentence Summary:** A robust approach for haplotype-resolved Hi-C reveals highly-structured homolog pairing in early stage *Drosophila* embryos.

---

Although chromosomes are organized within the nucleus into distinct territories, they nevertheless come into contact *in trans* (*1*). In diploid organisms, including mammals, such *trans* contacts may involve specific interactions, such as pairing, between the homologous maternal and paternal chromosomes. Although best known in meiotic cells (*2*), homolog pairing can also occur in somatic cells and influence gene expression via phenomena such as transvection (reviewed by (*2-7*)). In *Drosophila*, somatic pairing occurs throughout development, making this organism an ideal system for studying *trans* interactions (reviewed by (*2-7*)).

Homolog pairing was first noted in Dipteran insects, such as *Drosophila*, over 100 years ago (*8*). However, definitive demonstration of its extent and a molecular understanding of its structural basis have remained elusive. Recently, high-(bioRxiv (*9*)) and super-resolution (3D-SIM) (*10, 11*) imaging of several genomic loci has suggested that pairing involves the juxtaposition of chromosomal domains that nevertheless remain distinct, while simulations of pairing (*12*) via integration of the lamina-DamID data (*13*) and the Hi-C data (*14*) have predicted correlations between pairing and epigenetic domains. Furthermore, a transgene-based study of transvection has proposed a domain-based mode for pairing (bioRxiv (*15*)). These studies align with long-standing discussions about the establishment, maintenance, and stability of pairing as well as the relationship between pairing and transcription (reviewed by (*2-7, 16, 17*)). Nevertheless, the extent and the structure of pairing are largely unknown. Furthermore, it has been unclear whether pairing is uniform and truly genome-wide. For example, pairing may encompass many forms, ranging from a well-aligned juxtaposition of homologs (‘railroad track’), to a loose, ‘laissez-faire’ association of homologous regions, or even an apposition of highly disordered structures; for instance, pairing in mammalian systems can manifest as a nonrandom, approximate co-localization of homologous chromosomal regions (reviewed by (*4, 6*)). To address the structure of pairing, we have applied a haplotype-resolved form of Hi-C, which has previously been used to study *cis* interactions in mammalian systems (*18-29*) as well as *trans*-homolog interactions in yeast (*30, 31*). Here, we queried whether homologs in early *Drosophila* embryos are close enough to be captured by Hi-C and, if so, whether the captured *trans*-homolog signals are able to reveal the genome-wide *trans*-homolog architecture and its relationship to *cis*; as Hi-C ligates interacting genomic regions such that sequencing of the ends of the ligation products produces pairs of reads (read pairs) that identify the interacting genomic regions, it can be used to identify interactions between maternal and paternal homologs.

Strikingly, we discover that homolog pairing can be detected genome-wide by Hi-C, that it is extensive along entire lengths of chromosomes and highly structured, including *trans*-homolog domains, domain boundaries, and interaction peaks. We reveal the coincidence between *trans*-homolog features and the positions and sizes of analogous *cis* features and then consider our findings in light of three properties of homolog pairing: precision, proximity, and continuity. These findings are further pursued in our companion paper, which presents an in-depth analysis of the structure of pairing in a newly generated hybrid cell line (AlHaj Abed, Erceg, Goloborodko *et al.* bioRxiv (*32*)). In the current study, we focused our attention on the early embryo, uncovering further that *trans*-homolog organization correlates with the opening of chromatin during zygotic genome activation. Finally, we compare the extraordinary extent of homolog pairing in *Drosophila* embryos to *trans*-homolog contacts in mammalian embryos.

We focused our Hi-C analysis on the window of *Drosophila* embryogenesis extending from 2 to 4 hours after egg laying (AEL), when zygotic gene transcription is activated and the embryo transitions from a syncytial blastoderm into a multicellular state (*33, 34*). It is during this window that maternal and paternal homologs come together for the first time (*35-37*), and the rapid early cleavage cycles give way to the longer 13^th^ and 14^th^ cycles (*33*), thus minimizing the contribution of mitotic genome organization to our Hi-C signal (*38*). To generate the embryos for our study, we mated two divergent *Drosophila* Genetic Reference Panel lines, 057 and 439 (Fig. 1, A and B), as a high density of single nucleotide variants (SNVs) is required for assaying homolog pairing via genomic methods. Indeed, strains 057 and 439 differ with an average SNV frequency of ~5.5 per kilobase (kb), except on chromosome 4 (1.0 SNV/kb) (*39*). This frequency was adjusted to ~5.1 SNVs/kb (0.01 for chr4) when we re-sequenced both lines and reconstructed an F1 diploid genome using only high-confidence homozygous SNVs (fig. S1 and table S1, see supplementary materials and methods); this diploid genome assembly ensured the most stringent haplotype-resolved mapping of Hi-C products. Importantly, we confirmed that hybrid embryos achieved levels of homolog pairing consistent with those observed in other studies (*35-37, 40, 41*) using fluorescent *in situ* hybridization (FISH) to assess pairing (Fig. 1C) at heterochromatic (16.0%±5.7 to 62.4%±3.6) and euchromatic (2.0%±1.5 to 22.0%±2.4) loci across different chromosomes (Fig. 1, D and E). In addition, pairing increased to expected levels in 057/439 embryos as they aged (fig. S2A).

**Fig. 1.**
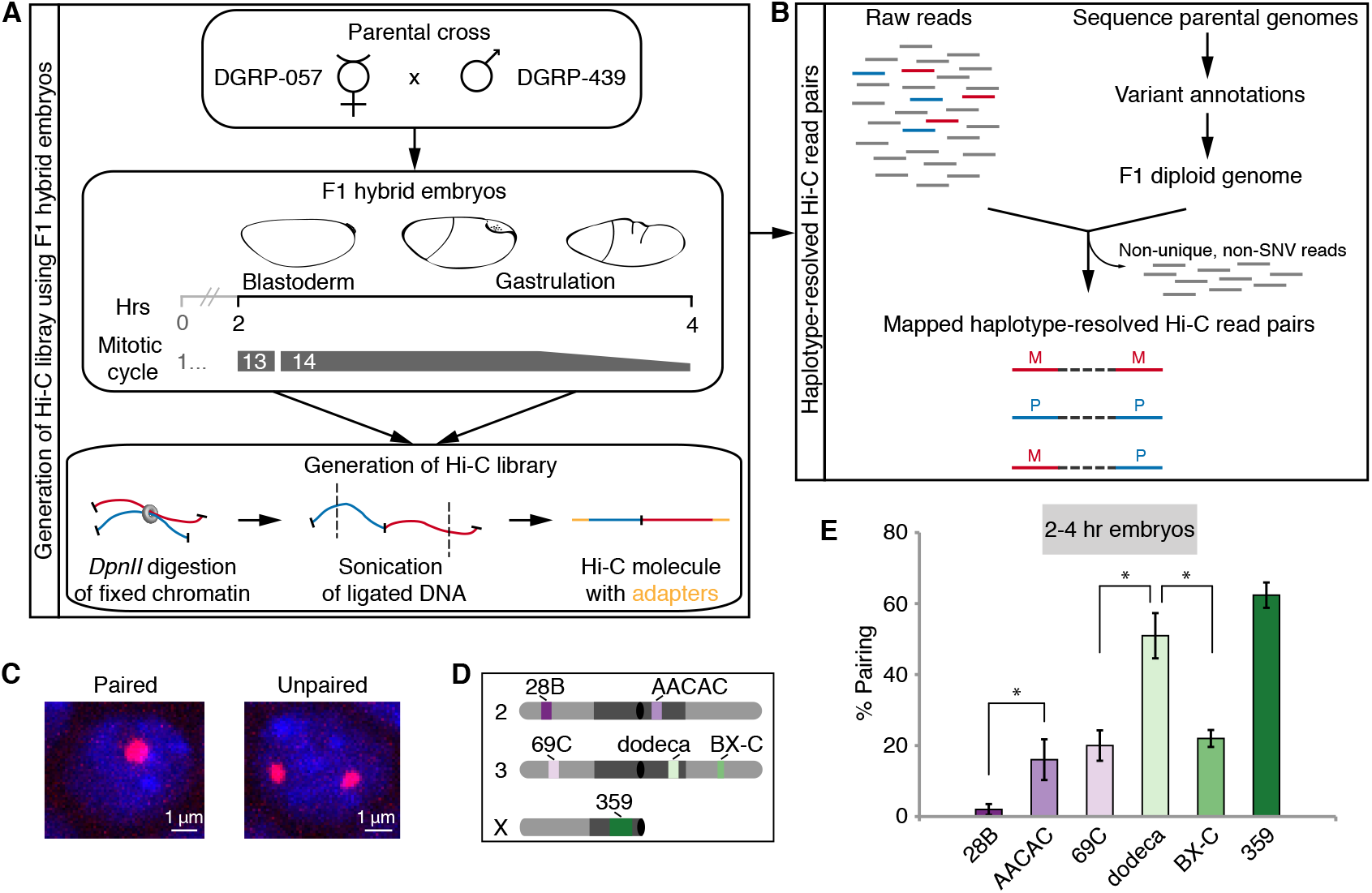
Haplotype-resolved Hi-C approach in F1 hybrid embryos that exhibit homolog pairing. (**A**) Generation of Hi-C libraries using 2-4 hr hybrid embryos and (**B**) haplotype-resolved mapping. (**C**) Homolog pairing as assessed by FISH. Nuclei are considered paired nuclei when FISH signals are ≤0.8 μm (center-to-center distance) apart. (**D**) Location of FISH targets (heterochromatin, dark grey; euchromatin, light grey; centromere, black circle). (**E**) Percentage of nuclei showing paired loci in 2-4 hr embryos (error bars, standard deviation of at least 3 replicates; n ≥100 nuclei/replicate; *, P<0.0001, Fisher’s two-tailed exact).

We performed Hi-C and recovered 513 million Hi-C products, of which ~56% provided unique read pairs that could be mapped to the reference genome (table S2). When assembled into a map and analyzed at 1 kb resolution without regard to parental origin of the mapped fragments, these reads revealed features that are routinely seen in non-haplotype-resolved Hi-C maps (*38, 42, 43*) – a central *cis* diagonal representing short-distance interactions in *cis* (*cis* read pairs) as well as signatures for compartments, domains, and interaction peaks – thus confirming the quality of our Hi-C data (fig. S2B). Presence of these interphase hallmarks indicates that the vast majority of the cells are in interphase rather than mitosis, where such features are absent (*38*).

Using our newly generated haplotype-resolved F1 diploid genome to determine the parental origin of our reads, we found that 5.8% of all mappable read pairs appeared to represent *trans* (*trans*-homolog as well as *trans*-heterolog) contacts and that, of these, 36.1% indicated contacts between homologous chromosomes (*trans*-homolog read pairs) (table S3). Encouraged by these numbers, we next determined the quality of the purported *trans*-homolog reads. For example, errors in reconstruction of the F1 diploid genome or sequencing of the Hi-C products can lead to misassignment of *cis* read pairs as *trans*-homolog read pairs (Fig. 2A) and, given the greater number of *cis* as *versus trans*-homolog read pairs, even a small rate of misassignment of *cis* read pairs could be confounding. Thus, to assess the potential magnitude of homolog misassignment, we sought a signature for it in our Hi-C dataset, focusing on read pairs where the separation along the genome (genomic separation) of the two reads of a read pair is <1 kb (Fig. 2B, shaded area). We chose to examine these read pairs because they are enriched in non-informative byproducts of the Hi-C protocol (*44*), the most abundant being unligated pieces of DNA, known as dangling ends (Fig. 2B, hatched area and fig. S3, see supplementary materials and methods). Dangling ends have genomic separations of <1 kb, are “inwardly” orientated, and 100-1,000 fold more abundant than read pairs of other orientations at the same separations (fig. S4A) and, since they are exclusively *cis* read pairs, enrichment of such pairs among *trans*-homolog read pairs can only arise via homolog misassignment (HM in figures and supplementary materials and methods). As such, the enrichment of short-distance (<1 kb) *trans*-homolog pairs can serve as a signature of homolog misassignment and be used to assess the overall quality of our homolog assignment (supplementary materials and methods). Consistent with this hypothesis, this signature of homolog misassignment almost disappears when we increased the stringency of our mapping by requiring at least 2 SNVs per read (Fig. 2C and fig. S4, B and C, probability of homolog misassignment P_HM_=0.01%). In contrast, homolog misassignment increased when we used the less accurate DGRP SNV annotations (fig. S4D, P_HM_=2.40%). This approach allowed us to estimate the probability of homolog misassignment as only ~0.17% when requiring at least 1 SNV per read (Fig. 2D, see supplementary materials and methods). Finally, this analysis indicated that homolog misassignment introduced less than 5% of erroneous *trans*-homolog contacts at separations above 1 kb (Fig. 2E). Having gained confidence that contamination contributes only a minor portion of purported *trans*-homolog read pairs, we observed that *trans*-homolog contacts at genomic separations of 1 kb are >100 fold more frequent than those at genomic separations of >1 Mb, and >500 fold more frequent than contacts between regions on different chromosomes (*trans*-heterolog, Fig. 2B). Thus, *trans*-homolog contacts constitute a major fraction of *trans*-chromosomal interactions.

**Fig. 2.**
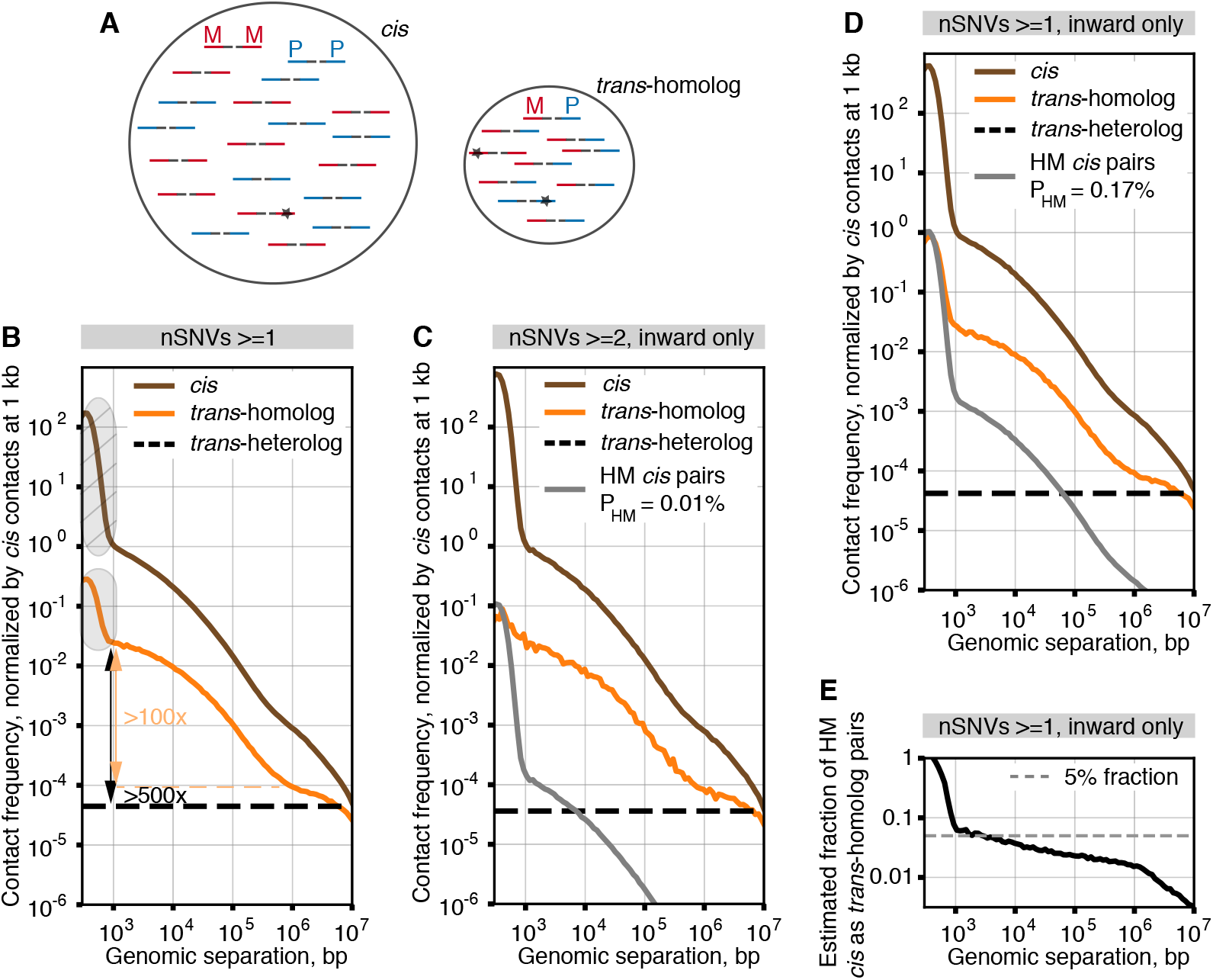
Strategy for robust distinction of *trans*-homolog from *cis* contacts. (**A**) Homolog misassignment can arise from sequencing errors, false SNVs, or sample heterogeneity (lines, Hi-C molecules; red, maternal fragment (M); blue, paternal fragment (P); stars, errors). (**B**) Contact frequency plotted against genome separation using ≤1 SNV per read. Arrows, change in contact frequency between short-range *trans*-homolog contacts and either long-range (> 1Mb) *trans*-homolog (orange) or *trans*-heterolog (black) contacts. Shaded area over the *trans*-homolog curve, enrichment of contact frequency due to homolog-misassigned “dangling ends” (shaded area over *cis* curve). (**C**) Contact frequency for inward only read pairs with at least 2 SNVs per read, with no sequence mismatches allowed (P_HM_=0.01%). (**D**) Contact frequency for inward only read pairs with at least 1 SNV per read (P_HM_=0.17%). (**E**) Fraction of homolog-misassigned *cis* contacts among *trans*- homolog pairs as a function of genomic separation. (**B** to **D**) Contact frequencies for chromosomes 2 and 3 normalized by *cis* contact frequency at 1 kb. Dashed black line, average *trans-*heterolog contact frequency; HM, homolog misassignment; P_Hm_, probability of homolog misassignment.

We next generated a Hi-C map using ≥1 SNV per read and filtering our data to exclude contacts below 3 kb of genomic separation in order to substantially reduce contamination by Hi-C byproducts. The resulting map was remarkably different from the non-haplotype-resolved Hi-C map (fig. S2B). In addition to displaying the expected central *cis* diagonal, it revealed prominent “*trans*-homolog” diagonals that extended across the entire length of each chromosome (Fig. 3A, black arrows). This observation demonstrated, first, that homolog pairing in *Drosophila* does indeed bring homologs close enough together to be detectable by Hi-C and, second, that this degree of homolog proximity occurs genome-wide.

**Fig. 3.**
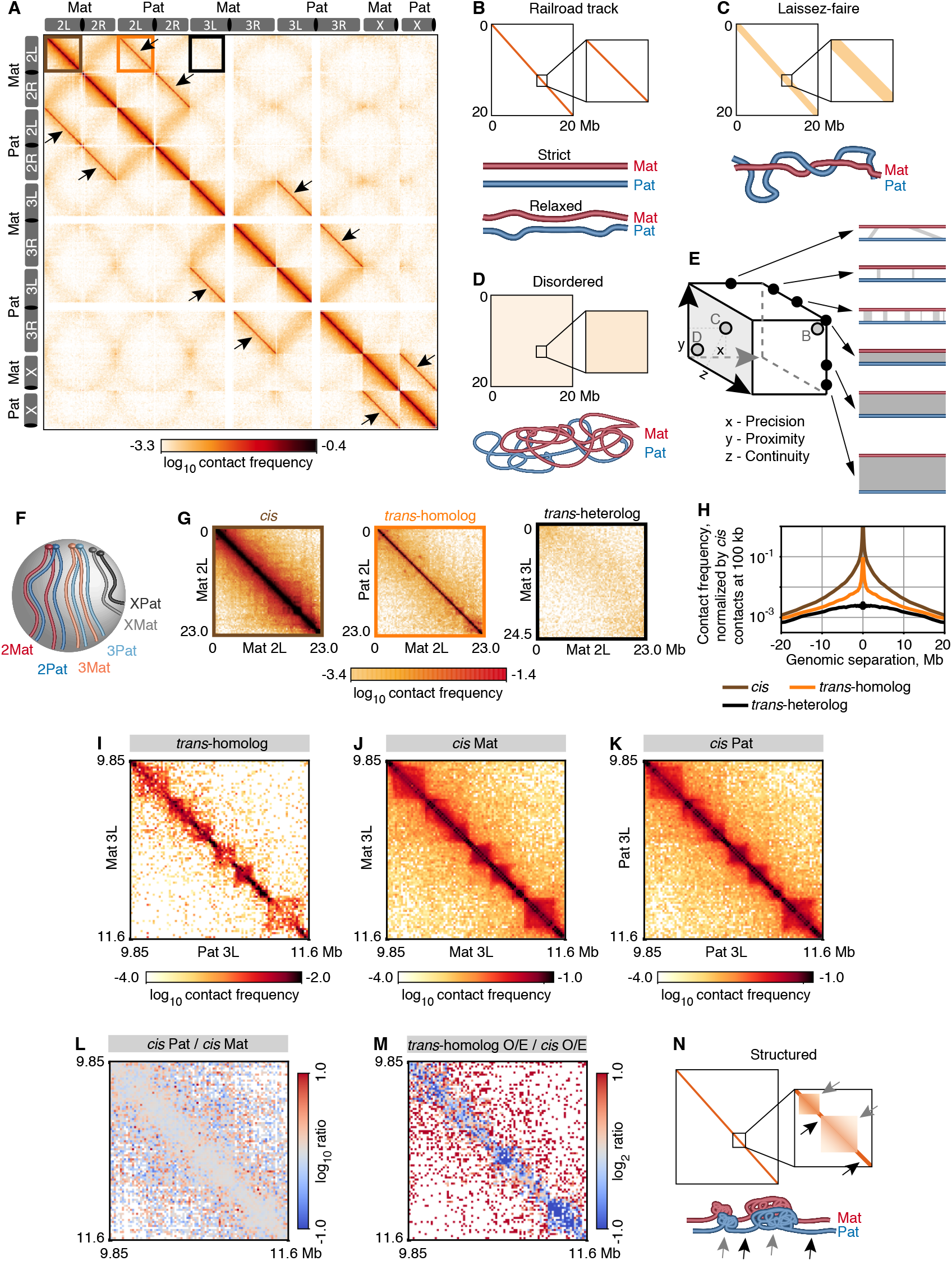
Genome-wide, multi-layered, and highly structured homolog pairing resembles features of *cis*-organization. (**A**) Haplotype-resolved Hi-C map of F1 hybrid embryos. Arrows, *trans*-homolog diagonals. Homologs juxtaposed in a (**B**) ‘railroad track’ fashion, (**C**) a loose, ‘laissez-faire’ mode, or (**D**) a highly disordered structure. (**E**) Homolog pairing may encompass a range of structures defined by precision, proximity, and continuity. (**F**) Rabl positioning of chromosomes. The (**G**) zoomed-in maps of chromosomal arms (color coded boxes in A; L, left; R, right) and (**H**) respective contact frequencies as a function of genomic distance (see supplementary materials and methods). (**I**) *trans*- homolog, (**J**) *cis* maternal, and (**K**) *cis* paternal maps of matching regions on chr3L. (**L**) Ratio of *cis* Pat/*cis* Mat Hi-C maps indicates the *cis* contact patterns of two homologs are highly consistent. (**M**) The ratio of *trans*-homolog/average *cis* maps suggests that pairing resembles *cis* contacts, albeit with lower interactions in some regions (dark blue). (**N**) Homolog pairing displays highly structured *trans*-domains (black) and *trans*-boundaries (grey), reflecting *cis*-organization of homologs.

The *trans*-homolog diagonals also suggest that homologs are relatively well-aligned when coming into contact, consistent with a ‘railroad track’ structure (Fig. 3B). This indication, however, does not preclude less aligned forms of pairing, such as the paired domains observed at a number of genomic regions by Szabo *et al.* (*11*). For example, Figure 2B also shows that extensive *trans*-homolog interactions may occur between non-allelic regions corresponding to genomic separations of hundreds of kilobases or more, albeit at much reduced frequencies. We would expect such interactions to appear as signals at varying distances off the *trans*-homolog diagonals, leaving open the possibility of other structures, such as laissez-faire (Fig. 3C) and highly disordered pairing (Fig. 3D).

That pairing may assume different forms in *Drosophila* has long been considered (reviewed by (*2-7*)), with recent studies suggesting that the state of pairing is a fine balance between pairing and anti-pairing factors (reviewed by (*6*)). Here, we propose a framework in which the different aspects of pairing can be placed along the continuum of three axes representing i) the precision with which homologous regions are aligned (x axis), ii) the proximity with which homologs are held together (y axis), and iii) the continuity or degree to which a particular state of pairing extends uninterrupted (z axis), with the continuum along any axis potentially reflecting different forms of pairing and/or the maturation (or degradation) of pairing over time (Fig. 3E). In this context, maximum values along all three axes would produce railroad track pairing, intermediate values would approximate a laissez-faire mode of pairing, and minimum values would result in extreme disorder. This framework can also be used to describe *trans* interactions between heterologous chromosomes, particularly prominent examples being those arising from the Rabl polarization of centromeres and telomeres, appearing in some Hi-C maps, including ours, as contacts along non-homologous arms (*38, 42, 43, 45*); the Rabl configuration may facilitate homolog pairing by reducing search space within the nucleus (Fig. 3F, (*35-37*)). Note, however, that *trans*-homolog contacts were tighter and/or more frequent than *trans*-heterolog contacts (Fig. 3, G and H), emphasizing the prevalence of homolog pairing in *Drosophila*.

We next asked whether our haplotype-resolved maps could further clarify the molecular structure of paired homologs as well as elucidate how *trans*-homolog interactions are integrated with the *cis* interactions that shape the 3D organization of the genome. Remarkably, we observed *trans*-homolog domains, domain boundaries, and interaction peaks resembling analogous features in *cis* maps (Fig. 3, I to K and fig. S5, A to C). In fact, 54% of the boundaries of *trans*-homolog domains overlapped a boundary of a *cis* domain (averaged over two homologs, fig. S5D). While this value is lower than the upper bound of 72% for *trans*-homolog boundaries shared between replicate haplotype-resolved Hi-C datasets, it is nevertheless significantly above the 35% overlap expected at random (P<10^-10^). Note that, as we obtained 92.3% overlap of domain boundaries between replicates of non-haplotype-resolved Hi-C datasets, it is likely that the overlap values for haplotype-resolved datasets reflect the relatively low number of haplotype-resolved reads we obtained.

The similarity between *trans*-homolog and *cis* contact maps is further highlighted through comparisons with maternal- and paternal-specific Hi-C domains, which have been observed in other systems to be concordant (*21*); in our dataset, we observed 73.9% concordance between *cis* domain boundaries of homologous chromosomes (fig. S5D; also Fig. 3L and fig. S5E), wherein concordance between replicates was 73.2% (see supplementary materials and methods). *Trans-*homolog maps do, however, differ from some *cis*-defined features (Fig. 3M and fig. S5F, dark blue), and we pursue this observation in our companion paper (Alhaj Abed, Erceg, Goloborodko *et al.* bioRxiv (*32*)). Taken together, our findings indicate that homolog pairing goes far beyond simple genome-wide alignments of homologous pairs and includes surprisingly ‘structured’ *trans*-homolog domains, boundaries, and specific interaction peaks that frequently correspond to *cis* features (Fig. 3N).

Spurred on by the rich history of transvection (reviewed by (*3-7*)) we asked how broadly pairing might be correlated the binding of major transcription factors in the early *Drosophila* embryo. We focused our attention on arguably the four most prominently studied maternally-contributed factors, involved in zygotic genome activation, in particular key developmental events such as the opening of chromatin, transcription, and embryonic pattern formation: the pioneer factor Zelda (Zld) (*46, 47*), which mediates early chromatin accessibility; Bicoid (Bcd) (*48*) and Dorsal (Dl) (*49, 50*), which are involved in patterning the anterior-posterior and dorsal-ventral axis, respectively, and GAGA factor (GAF) (*51, 52*), which may facilitate transcription in later stages. We correlated the binding of these factors, as revealed by ChIP-seq datasets (*48, 50, 53*), to local variations in the degree of pairing as characterized by the ‘pairing score (PS)’, defined as the log_2_ *trans*-homolog contact frequency within a 28 kb window along the diagonal of Hi-C maps (see supplementary materials and methods). Excitingly, regions of elevated PS were significantly enriched for Zld and Bcd (P<10^-10^, Fig. 4, A and B). Interestingly, 25% of the top Zld-occupied regions have high PS (P<10^-10^, Fig. 4C). In contrast, neither GAF nor Dl were positively correlated with the PS (P = 0.139, P=0.029, respectively; Fig. 4B).

**Fig. 4.**
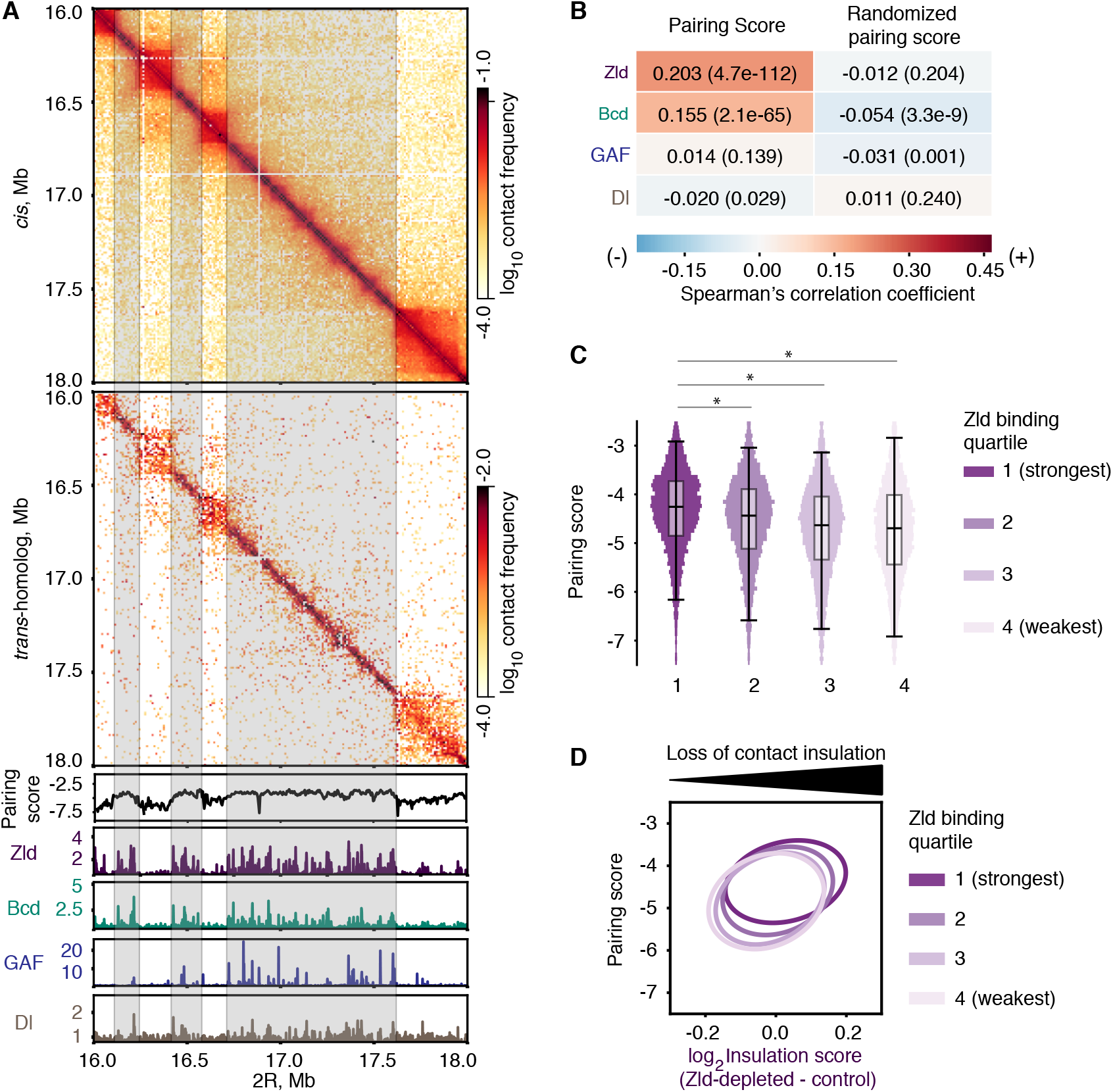
Homolog pairing is related to Zld-mediated opening of chromatin. (**A**) *Cis* (upper panel) and *trans*-homolog (second panel from the top) contact maps of a 2 Mb region (2R:16-18 Mb). Lower panels, pairing score (PS) calculated using a 28 kb window at 4 kb resolution (black), ChIP-seq profiles of Zld (dark purple) (*50*), Bcd (green) (*48*), GAF (blue) (*53*), and Dl (brown) (*50*). Grey boxes, regions of elevated PS in boundaries. (**B**) Correlation analyses between the PS and the binding profiles of Zld, Bcd, GAF, and Dl as determined using pairwise Spearman correlation. Control, randomized 200 kb chunks of the PS. Spearman correlation coefficients indicated in each box and by a heatmap; P-values in parentheses. (**C**) 25% of the strongest Zld binding coincides significantly with high PS regions, compared to the remaining Zld binding (*P<10^-10^, Mood’s median test). (**D**) PS *vs* loss of contact insulation upon Zld depletion, grouped by Zld binding strength quartile. Ovals represent the contours of 2D-Gaussian approximations, at a distance of 2 standard deviations from the mean (the dot). Regions with stronger Zld binding show both high PS and increased loss of contact insulation upon Zld depletion.

As previous studies suggested that regions with high Zld occupancy were more dependent on Zld for establishing domain boundaries (*38*), we asked whether Zld may play a role in domains that are established via *trans*-homolog interactions. Three observations supported our hypothesis. First, visual inspection of our Hi-C revealed that *trans*-homolog domain boundaries are associated with high Zld occupancy and, moreover, are associated with values of PS that are higher than those of the interior of domains (Fig. 4A and fig. S6A, P<10^-10^). Second, using Hi-C data from Zld-depleted embryos (*38*), we observed a decrease of boundary strength at regions that would otherwise be associated with high PS values and strong Zld binding (Fig. 4D and fig. S6B, P<10^-10^). Finally, we found that Zld depletion resulted in reduced pairing at region otherwise associated with high Zld occupancy, but not regions associated with low Zld binding (P = 2.56×10^-3^, fig. S6, C and D). Thus, Zld-mediated chromatin accessibility may be related to the establishment of homolog pairing in the early embryo (Fig. 3N).

One of the most exciting developments in recent years is the growing indication that homolog pairing is not special to *Drosophila*, that it may be a general property in many organisms, including mammals (reviewed by (*4, 6*)). Thus, using our computational approach for detecting homolog pairing through Hi-C, we assessed the state of pairing in mammalian embryos. As pairing in mammals is generally transient and localized, for example, during DNA repair, V(D)J recombination, imprinting, or X-inactivation (reviewed by (*4, 6*)), these analyses would also serve as a control for observations in *Drosophila*, where pairing is far more extensive. In particular, we considered two deeply sequenced Hi-C datasets from Du *et al.* (*25*) and Ke *et al.* (*26*) representing gametes and F1 hybrid mouse embryos bearing a high frequency of SNVs (1 autosomal SNV per ~130 bp or ~490 bp, respectively). Reasoning that the haploid nature of sperm and oocytes precludes homolog pairing, we focused first on these cell types to determine the level of stringency that would be necessary to remove any apparent *trans*-homolog read pair that would necessarily have resulted from homolog misassignment (fig. S7A). That level of stringency was ≥2 SNVs per read (fig. S7B and table S4, see supplementary materials and methods). Using this protocol, we turned to the datasets representing embryos at the 8-cell stage or earlier and observed no clear signal for *trans*-homolog contacts (figs. S8 and S9). These observations aligned with previous indications that the maternal and paternal genomes are spatially segregated in zygotes (*54, 55*) and remain so to a lesser extent as late as the 8-cell stage ((*25, 26*); fig. S8).

In contrast, the signal for *trans-*homolog contacts was significantly higher than expected from homolog misassignment alone across a range of genomic separations in the older E7.5 embryos of Ke *et al.* (fig. S8). While this weak signal is consistent with a low level of homolog pairing at genomic separations of over 10 kb to almost 10 Mb, it could also reflect the clustering of SNVs and/or Hi-C biases (see (*44*)) that could not be completely eliminated due to sparsity of the data (table S4). This underscores the extraordinary degree of somatic homolog pairing in *Drosophila*, even as they leave open the possibility of pairing in early mammalian embryos.

To conclude, our study addressed the fundamental nature of chromosome pairing in *Drosophila.* We developed a robust approach for conducting haplotype-resolved Hi-C studies, wherein interactions between homologous chromosomes are carefully vetted for homolog misassignment. Using this approach, we obtained, for the first time, a genome-wide high-resolution view of homolog pairing during early *Drosophila* development. Our data revealed genome-wide juxtaposition of homologs along their entire lengths as well as a multi-layered organization of *trans*-homolog domains, domain boundaries, and interaction peaks. We also observed concordance of *trans*-homolog and *cis-* features, arguing that *cis* and *trans* interactions are structurally coordinated (pursued further by AlHaj Abed, Erceg, Goloborodko *et al*. bioRxiv (*32*)). This striking correspondence of *trans*-homolog and *cis* domain boundaries and interaction peaks may suggest similar mechanisms of their formation. For instance, SMC complexes implicated in loop extrusion (*56*), formation of *cis* domains, and cohesion of sister chromatids in mammals may play a role in pairing and formation of *trans*-homolog domains in *Drosophila*.

Moreover, by layering *trans*-homolog contacts on models of genome organization that have relied primarily, if not solely, on *cis* contacts, our findings highlight how haplotype-resolved Hi-C can clarify paradigms of genome organization and function. Indeed, pairing may underlie some structural and regulatory differences between the 3D architecture of the *Drosophila* genome architecture and that of other organisms that lack pairing. Our observations of a correlation between pairing and Zelda-mediated chromatin accessibility during zygotic genome activation champion a relationship between *trans*-homolog genome organization and key developmental decisions, aligning well with the growing recognition that pairing can play a potent role in gene regulation, even at some loci in mammals (reviewed by (*4, 6*)). While we found no definitive evidence of extensive genome-wide pairing in available embryonic haplotype-resolved mouse data, our study nevertheless leaves open the possibility of transient and/or locus-specific paring. Finally, we believe that the sensitivity of our Hi-C approach has potential usefulness also for the analysis of *trans* contacts between heterologous chromosomes. In the framework of precision (x), proximity (y), and continuity (z), such *trans*-heterolog contacts would fall in the plane of x = 0. Of course, when this framework is extended to encompass the dimension of time, it should then be able to capture the dynamics of *trans*-homolog and *trans*-heterolog contacts through cell division, development, aging, and disease.

## Acknowledgments

We are grateful to the Wu and Mirny laboratories, participants of the Annual Northeast Regional Chromosome Pairing Conferences, the Lieberman Aiden laboratory, 4DN DCIC, in particular Giancarlo Bonora from the Noble laboratory, Brian J. Beliveau, Guillaume J. Filion, Mirko Francesconi, Ben Lehner, M. Jordan Rowley, and the Cavalli laboratory, especially Frédéric Bantignies, for discussions, the Furlong laboratory for Oregon-R fly collection, the Rushlow laboratory for *UAS-shRNA-zld* line, the TUCF Genomics Sequencing Core Facility, the Microscopy Resources on North Quad (MicRoN), and the Bloomington *Drosophila* Stock Center for *Drosophila* stocks.

## Funding

This work was supported by an EMBO Long-Term Fellowship (ALTF 186-2014) to J.E., a William Randolph Hearst Award to R.B.M., and awards from NIH/NCI (Ruth L. Kirschstein NRSA, F32CA157188) to E.F.J., NIH/NHGRI (R01 HG003143) to J.D. (Howard Hughes Medical Institute investigator), NIH/NIGMS (R01 GM114190) to L.A.M., and NIH/NIGMS (RO1GM123289, DP1GM106412, R01HD091797) and HMS to C.-t.W. J.D. and L.A.M. acknowledge support from the National Institutes of Health Common Fund 4D Nucleome Program (U54 DK107980).

## Author contributions

J.E., J.A.A., and C.-t.W. designed the experiments. A.G. designed Hi-C computational analyses with input from J.E., J.A.A., L.A.M. and C.-t.W. J.D. and B.R.L. provided input in experimental design and data analysis. Experimental data were generated by J.E. and J.A.A., with the help of S.C.N., E.F.J., T.N.S., M.A.H. for sorting males and virgin females, and G.N. for Oligopaints design. J.E. and R.B.M. selected the parental lines. J.E., J.A.A., A.G., B.R.L., G.F., N.A., and M.I. performed computational analyses of the Hi-C data. J.E. performed FISH data analysis and Zld depletion experiments. J.E., J.A.A., A.G., G.F., L.A.M., and C.-t.W. interpreted the data. J.E., J.A.A., A.G., L.A.M., and C.-t.W. wrote the paper with input from the other authors.

## Competing interests

Authors declare no competing interests.

## Data and materials availability

Raw sequencing data and extracted Hi-C contacts have been deposited in the Gene Expression Omnibus (GEO) repository under accession number GSE121255.

## Materials and Methods

### Selection of parental *Drosophila* lines

To be able to distinguish homologs in sequencing data, we selected two parental lines with a high number of SNVs from *Drosophila melanogaster* Genetic Reference Panel (DGRP), which contains *Drosophila* lines inbred over 20 generations (*39*). To ensure high quality and accuracy of variant calling, we used only those lines whose sequence coverage with Illumina technology was at least 20x. Variant annotation for each fly line with at least 20x sequence coverage was extracted from Freeze 2 Release (https://www.hgsc.bcm.edu/arthropods/drosophila-genetic-reference-panel; freeze2.bins.vcf.gz), and a count of the number of SNVs against reference genome (VCFTtools, vcf –stats function) (*57*) was annotated. We proceeded with the top 5 and bottom 5 ranked *Drosophila* lines by filtering their variant annotations to keep only homozygous SNVs, and then performed pairwise comparison (vcftools --vcf DGRP-x_homSNV.vcf --diff DGRP-y_homSNVs.vcf). Upon inspection of the output file from the pairwise comparison (out.diff.site_in_files), which contained information on both common and unique homozygous SNVs between two lines, we selected two parental lines, DGRP-057 and DGRP-439, that differed by most homozygous, fixed SNVs.

### Genomic DNA isolation and library preparation

20 adult flies from each parental line (the Bloomington stock numbers 29652 and 29658) were homogenized with motor (Kimble-Kontes Pellet Pestle Cordeless Motor) and pestle (Kimble-Kontes) on ice. Genomic DNA was isolated using DNeasy Blood & Tissue Kit (Qiagen) according to manufacturers’ recommendations. The integrity of the isolated genomic DNA (*i.e.* no degradation present) was verified by running a 1% agarose gel. Genomic DNA libraries were generated using Illumina TruSeq Nano DNA Library Preparation kit, and were 150 bp paired-end sequenced at the TUCF Genomics Facility using Illumina HiSeq2500.

### Collection and fixation of the F1 hybrid embryos

The inbred DGRP-057 (maternal) and 439 (paternal) lines were mated to obtain the F1 hybrid embryos. To ensure that accurate parental genotypes were crossed, around 28,000 males and 28,000 virgin females were manually sorted. The F1 hybrid embryos were collected and fixed after 3 pre-lays followed by aging for 2-4 hr time-point at 25°C as previously described (*58*). To validate that the embryos were of the correct stage, and did not contain contaminants from older embryos, a small aliquot (~100 embryos) per collection was set aside, devitellinised, and stored in methanol at −20°C to verify developmental stages of collection under microscope. If even a single older embryo was noted, collection was discarded, as that embryo may contribute more nuclei than 2-4 hr embryo, depending on exact developmental difference between them. The remaining embryos from collections were snap-frozen in liquid nitrogen and stored at −80°C.

### *in situ* Hi-C on *Drosophila* F1 hybrid embryos

Hi-C protocol from Rao *et al.* (*19*) was adapted for the *Drosophila* embryos. 30 mg of snap-frozen fixed embryos were homogenized with pestle (Kimble-Kontes) in 500 μl of ice-cold lysis buffer, and incubated 30 min on ice. Upon cell lysis, the chromatin was digested using 500 U of DpnII restriction enzyme overnight at 37°C. The DpnII overhangs were filled in with biotin mix during 1.5 hr at 37°C, and the chromatin was ligated for 4 hr at 18°C. RNA was degraded by adding 5 μl of 10 mg/ml RNase A (ThermoFisher Scientific) followed by incubation for 30 min at 37°C. After degradation of proteins and crosslink reversal overnight at 68°C, DNA was ethanol precipitated. DNA shearing was performed to obtain around 400-600 bp fragments with Qsonica Instrument (Q800R) using the following parameters: 30 sec on/off, 15x cycles, 70% ampl. After size-selection to around 500 bp, biotin pull-down was performed with a minor modification. Namely, the size-selected DNA was eluted in 200 μl of 10 mM Tris-HCl pH 8.0, and was mixed with equal volume of 2x Binding Buffer (10 mM Tris-HCl pH 7.5, 1 mM EDTA, 2 M NaCl) containing pre-washed Dynabeads MyOne Streptavidin T1 beads (Life Technologies).

The library preparation was performed as previously described (*19*) with the following modifications. The adapter ligation was in 46 μl of 1x Quick ligation reaction buffer (NEB), 2 μl of DNA Quick ligase (NEB), and 2 μl of NEXTflex DNA Barcode Adapter (25 μM, NEXTflex DNA Barcodes −6, Bioo Scientific #514101). PCR reaction contained 23 μl of adaptor ligated DNA, 2 μl of NEXTflex Primer Mix (12.5 μM), and 25 μl NEBNext Q5 Hot Start HiFi PCR Master Mix (NEB), and was run using the program: 98°C, 30 sec, (98°C, 10 sec, 65°C, 75 sec) repeated 7 times, 65°C, 5 min, hold at 4°C. Purification of PCR products was performed by incubation for 15 min instead of 5 min after each addition of Agencourt AMPure beads. The final libraries were eluted in 15 μl of 10 mM Tris-HCl pH 8.0 after 15 min incubation at room temperature, and 15 min incubation on a magnet. The library quality was assessed using the High Sensitivity DNA assay on a 2100 Bioanalyzer system (Agilent Technologies). The libraries corresponding to two independent biological replicates were 150 bp paired-end sequenced at the TUCF Genomics Facility using Illumina HiSeq2500.

### FISH probes

Oligo probes for heterochromatin repeats at chr2 (AACAC), chr3 (dodeca) and chrX (359) were previously described (*59-61*). Probes were synthesized by Integrated DNA Technologies (IDT) with the following sequences and a fluorescent dye: AACAC (Cy3-AACACAACACAACACAACACAACACAACACAACAC), dodeca (FAM488-ACGGGACCAGTACGG), and 359 (Cy5-GGGATCGTTAGCACTGGTAATTAGCTGC).

The libraries for euchromatin probes 28B (680 kb), 69C (674 kb), and 89D-89E/BX-C (315 kb) were designed using Oligopaint technology (table S5) as previously described (*41, 62, 63*), and were amplified using T7 amplification (see section ‘Oligopaint probe synthesis’) with forward primers containing a site for secondary oligo annealing, and reverse primers containing a T7 promoter sequence (sequences are provided in table S6). DNA secondary oligos (*62*), which were used together with primary probes, had both 5’ and 3’ conjugated fluorophores (table S6).

### Oligopaint probe synthesis

Oligopaints probes were synthesized using T7 amplification followed by reverse transcription as previously described (*64, 65*). Oligopaints libraries were first amplified with Kapa Taq enzyme (Kapa Biosystems, 5 U/μl), and corresponding forward and reverse primers (without the site for secondary oligo annealing and the T7 promoter, *i.e.* without the underlined sequences from table S6) using linear PCR program as follows: 95°C, 5 min, (95°C, 30 sec, 58°C, 30 sec, 72°C, 15 sec) repeated 25 times, 72°C 5 min, hold at 4°C. Upon clean up with DNA Clean & Concentrator-5 (DCC-5) kit (Zymo Research), this PCR was followed by another bulk up PCR using the same program, but this time to introduce the site for secondary oligo annealing to the forward strand, and the T7 promoter to the reverse strand (primers used *with* the underlined sequences from table S6). The PCR product with added T7 promoter was cleaned up using DNA Clean & Concentrator-5 (DCC-5) kit (Zymo Research), and became a template in T7 reaction to produce excess RNA with HiScribe T7 High Yield RNA Synthesis Kit (NEB) and RNAseOUT (ThermoFisher Scientific) overnight at 37°C. RNA was then reverse transcribed into DNA with Maxima H Minus RT Transcriptase (ThermoFisher Scientific) in a 2 hr reaction at 50°C, followed by the inactivation of the RT enzyme at 85°C for 5 min. Subsequently, RNA was degraded via alkaline hydrolysis (0.5M EDTA and 1M NaOH in 1:1) for 10 min at 95°C, and ssDNA was purified with DNA Clean & Concentrator-5 (DCC-5) kit (Zymo Research), where DNA binding buffer was replaced with Oligo binding buffer (Zymo).

### FISH in embryos

FISH in *Drosophila* embryos was performed as previously described with modifications (*41, 62*). Embryos were dechorionated with 50% bleach for 2.5 min, washed with 0.1% Triton X-100 in PBS, and fixed for 30 min in 500 μl of fix (PBS containing 4% formaldehyde, 0.5% Nonidet P-40, and 50 mM EGTA), plus 500 μl of heptane. Upon replacing the aqueous phase with methanol, embryos were vigorously shaken for 2 min. Finally, embryos were triple washed with 100% methanol, and stored at −20°C in methanol.

Prior to FISH, embryos were gradually rehydrated in 2x SSCT (0.3 M NaCl, 0.03 M NaCitrate, 0.1% Tween-20). Subsequently, embryos were incubated for 10 min in 2x SSCT/20% formamide, followed by another 10 min in 2x SSCT/50% formamide, and then hybridized in a hybridization solution (2x SSCT, 50% formamide, 10% dextran sulphate, RNase) with primary Oligopaint probe set (200 pmol for heterochromatin and 300 pmol for euchromatin probes) for 30 min at 80°C, and then at 37°C overnight. After primary probe hybridization, two 30 min washes in 2x SSCT/50% formamide were performed at 37°C. In a case secondary probe was used, secondary hybridization was performed between those two washes for 30 min in 2x SSCT/50% formamide with 300 pmol of secondary probe at 37°C. The washes were continued in 2x SSCT/20% formamide for 10 min at RT, and with three rinses in 2x SSCT. The embryos were stained with Hoechst 33342 (Invitrogen), washed in 2x SSCT for 10 min at RT, quickly rinsed in 2X SSC, and mounted in SlowFade Gold antifade mountant (Invitrogen). Images were taken using a Zeiss LSM780 laser scanning confocal microscope with a 63x oil NA 1.40 lens at 1024×1024 resolution.

### Analysis of FISH signals

The analysis was performed by manually examining each section of Z-stack with the Zeiss ZEN Software. Distance in 3D space between two FISH signals was measured using the Ortho-distance function in the Zeiss ZEN Software. Two homologs were considered paired if 3D distance between centers of two FISH signals was ≤0.8 μm or there was only one FISH signal.

### Generation of Zld-depleted embryos

Zld-depleted embryos were obtained as previously described (*38, 50*). The *UAS-shRNA-zld* (*50*) virgin females were crossed to the maternal triple driver Gal4 (*MTD-Gal4*, the Bloomington stock number 31777) males. The F1 heterozygous females *UAS-shRNA-zld* /*MTD-Gal4* were then mated to their sibling males, and the F2 Zld-depleted embryos were collected, and fixed as described in the section ‘FISH in embryos’. The control embryos were obtained using the same crossing scheme just by replacing the *MTD-Gal4* with the wild-type Canton S males. To validate that embryos were of the correct genotype, a small aliquot (~500 embryos) was inspected under microscope for both Zld-depleted and control samples.

### The construction of F1 diploid *Drosophila* genome

We sequenced the two inbred parental *Drosophila* lines (DGRP-057 and DGRP-439) together with the hybrid PnM cell line from AlHaj Abed, Erceg, Goloborodko *et al.* bioRxiv (*32*) at the average coverage of 118, 117 and 396 reads per base pair, respectively (*66*). We then detected the sequence variation of these three libraries using *bcftools*. In summary, we obtained high-quality normalized sequence variants as following:

1. Trimmed low-quality sequences with ‘seqtk trimfq’, aligned whole genome paired-end reads against the reference dm3 genome using BWA mem, and removed aligned PCR duplicates with ‘samtools --rmdup’.
2. Piled alignments up along the reference genome with ‘bcftools pileup --min-MQ 20 --min-BQ 20’,
3. Called raw sequence variants from the pileups with ‘bcftools call’.
4. Normalized raw sequence variants with ‘bcftools norm’.
5. Selected only high-coverage high-quality normalized sequence variants using ‘bcftools filter INFO/DP > 80 & QUAL > 200 & (TYPE="SNV" | IDV > 1)’.

We then phased heterozygous PnM variants using ‘bcftools isec’. We picked high-confidence variants on the maternal autosomes by selecting heterozygous PnM variants that were present among maternal DGRP-057 variants and absent among paternal DGRP-439 sequence variants (both homo- and heterozygous); the high-confidence paternal variants phasing was selected in an opposite manner.

Since PnM is a male line, for the maternal copy of chrX in F1 embryos, we considered only homozygous high-quality variants detected in the PnM cell line. To reconstruct the consensus sequence of the paternal copy of chrX in F1 embryos, we kept only homozygous variants detected in the paternal DGRP-439 *Drosophila* line.

Finally, we reconstructed the sequence of the F1 embryos with ‘samtools consensus’, using (a) the homozygous autosomal PnM SNVs, (b) the heterozygous phased autosomal PnM SNVs and (c) the homozygous maternal chrX PnM SNVs as well as the homozygous SNVs detected in the paternal DGRP-439 *Drosophila* line.

### Hi-C data analysis

#### Mapping and parsing

We started with sequences of Hi-C molecules and trimmed low-quality base pairs at both ends of each side using the standard mode of *seqtk trimfq* v.1.2-r94 (https://github.com/lh3/seqtk). Then we mapped the trimmed sequences to the reference dm3 genome or the constructed dm3-based F1 diploid genome using *bwa mem* v.0.7.15 (*67*) with flags -SP.

We then extracted the coordinates of Hi-C contacts using the *pairtools parse* command line tool (https://github.com/mirnylab/pairtools). We only kept read pairs that mapped uniquely to one of the two homologous chromosomes and removed PCR duplicates using the standard mode of the *pairtools dedup* command line tool.

#### Contact frequency (P(s)) curves

We used unique Hi-C pairs to calculate the functions of contact frequency P(s) *vs* genomic separation s. We grouped genomic distances between 10 bp and 10 Mb into ranges of exponentially increasing widths, with 8 ranges per order of magnitude. For every range of separations, we found the number of observed *cis*- or *trans*-homolog interactions within this range of separations and divided it by the total number of all loci pairs separated by such distances.

We used a similar technique to quantify the degree of coalignment of chromosomal arms due to Rabl configuration (Fig. 3H). Specifically, the Rabl configuration increased the contact frequency between pairs of loci located at the same *relative* positions along different chromosomal arms. We characterized this tendency by calculating a P(s) on *trans*-heterolog maps, where we calculated the position mismatch as s=x_1_-x_2_/L_2_*L_1_, where x_1_ and x_2_ were the distances from each locus to the centromere and L_1_ and L_2_ were the lengths of the two chromosomal arms.

#### Estimating the rate of Hi-C byproducts

Each Hi-C dataset contains non-informative byproducts, *i.e.* sequenced DNA molecules that originate from a single DNA fragment, and thus do not carry any information on chromatin conformation (unlike informative Hi-C molecules that form via a ligation of two spatially proximal DNA fragments and capture a spatial contact) (fig. S3A to B). The two well-known types of Hi-C byproducts are: (i) unligated DNA fragments (termed “dangling ends”) and (ii) self-ligated DNA fragments (“self-circles”) (*44*).

“Dangling ends” are small pieces of intact unligated DNA that passed through all selection steps of the Hi-C protocol and ended up in the library of sequenced molecules (fig. S3B). “Dangling ends” appear as *cis* read pairs at s~100-1,000 bp (such molecules are left in the library after the size selection step of the Hi-C protocol), with the direction of the two reads of a read pair pointing towards each other along the reference genome.

“Self-circles” are pieces of DNA whose ends were ligated to each other and the resulting circular DNA had a break in another location, producing a linear piece of DNA (fig. S3B). “Self-circles” also appear in *cis*, but at longer separations, up to a few kb (this distance depends on the frequency of the restrictions sites along the genome) and the resulting paired read orientations point away from each other along the reference genome (*44*).

Because both of these types of byproducts were characterized by their narrow range of genomic separations and a specific mutual orientation of the paired end reads (fig. S3B), their abundance was estimated using P(s) curves for four possible combinations of the directionality of paired read ends (fig. S3C and fig. S4A). In such plots, “dangling ends” produced a characteristic 100x-1,000x enrichment of inward read pairs below 1 kb separation and “self-circles” produced a weaker enrichment of outward read pairs at separations up to 10 kb, though their amount and typical genomic separation seemed to vary greatly between different experimental protocols (fig. S3C and fig. S4A) (*44*).

During the work on this manuscript, we also discovered a third, previously undescribed type of a Hi-C byproduct. We found that the read pairs that aligned to the same genome strand (*i.e.* same-strand read pairs) were enriched at s<500 bp. Upon closer examination, we found that such short-distance same-strand read pairs shared a similar unique pattern: while one of the two reads were typically fully aligned to some locus on the genome, the other read would split into two fractions, which would align to different locations. The inner, 3’ fraction mapped in *cis*, downstream and opposite to the read on the first side (as in a “dangling end”); while the outer, 5’ fraction overlapped the inner fraction, but went into the opposite direction (fig. S3D). This non-trivial alignment pattern suggested a scenario, where, during some stage of the Hi-C protocol, a DNA molecule formed a hairpin and was extended using itself as a template (fig. S3E). During the work on this manuscript, this type of a Hi-C byproduct was independently reported and thoroughly characterized in (*68*), who dubbed it as “hairpin loops”. Another, older study reported formation of similar hairpin loops in whole-genome sequencing of ancient DNA (*69*).

The mechanism by which such “hairpin loops” form remains to be firmly established. Golloshi *et al.* (*68*) proposed that these “hairpin loops” were formed during the end repair stage of the Hi-C protocol via self-annealing of a pre-existing microhomology region followed by an elongation by the T4 polymerase.

If this hypothesis was true, then “hairpin loops” should form with equal probability on all Hi-C molecules, regardless of the orientation and *relative* location of the ligated fragments, as well as on “dangling ends” and “self-circles”. The enrichment in short distance same-strand read pairs would then be most likely due to formation of “hairpin loops” on “dangling ends”, which is a highly enriched substrate. Importantly, we can still identify the ligated fragments even in Hi-C molecules with “hairpin loops”. First, the extra base pairs introduced by a “hairpin loop” map to the same location as the substrate (albeit at the opposite direction) and their alignment can serve as a proxy for the location of the substrate. Second, provided a sufficient long read length, we can simply ignore these extra pairs and instead align the inner fraction of the Hi-C molecule, which served as a substrate for the “hairpin loop”. Thus, “hairpin loops” represent the most benign type of a Hi-C byproduct and their presence does not significantly reduce the number and identity of contacts extracted from a Hi-C library.

We produced an upper estimate of the amount of byproduct types in our sample, assuming that the true P(s) followed that of the same-strand read pairs at s>1 kb, and was constant at s<1 kb. Using this assumption, we found that “dangling ends” constituted 39.5% of mapped and phased reads in our Hi-C librarys, “self-circles” were 0.3%, and “hairpin loops” formed from “dangling ends” made up 2.9% of the library.

#### Binning and balancing of Hi-C data

We aggregated unique Hi-C pairs into genomic bins of 1 kb and larger, using the cooler package (https://github.com/mirnylab/cooler). Low signal bins were excluded prior to balancing using the MADmax filter: we removed all bins, whose coverage was 7 genome-wide median deviations below the median bin coverage. Additionally, we removed all *cis*- and *trans*-homolog contacts at separations below 3 kb, since these reads pairs were dominated by non-informative Hi-C artifacts, unligated DNA fragments and ligation sites formed via self-circularization of DNA fragments (*70*). We then balanced the obtained contact matrices via iterative correction (IC), *i.e.* equalized the sum of contacts in every row/column to 1.0.

We calculate observed/expected contact frequency maps (often abbreviated as O/E CF) in *cis* and *trans*-homolog by dividing each diagonal of an IC contact map by its chromosome-wide average value over non-filtered genomic bins.

### Estimating the level of homolog misassignment

#### The definition of homolog misassignment

When we aligned a read to a diploid genome, we assigned to it three bits of information: which pair of homologous chromosomes this read originated from; the allele, *i.e.* which of the two homologs it originated from, and its location along the chromosome. Importantly, we could assign the allele (or, the homolog) to a read only if it overlapped some sequence that is unique to one homolog, *i.e.* if it overlapped a SNV or an indel; otherwise, it was impossible to tell which of the two homologs the read originated from.

Even in the *Drosophila* and mouse models with the highest density of SNVs, the sequences of the two homologous chromosomes were still highly similar and contained only one SNV per 100-200 bp. Because of that, the majority of read pairs (150 bp) lacked allele assignment on one or both sides; and the allele assignment for the rest of pairs was based on one or a few SNVs, and thus was prone to errors.

#### Sources of homolog misassignment

The errors in allele assignment (below, homolog misassignment, HM) occur for two reasons: (A) due to an error in the sequence of the read, or (B) due to an error in the sequence of the genome. Below we discussed how such errors lead to HM. Also, note that we only discussed the case where diploid genome sequences only contained SNVs and did not analyze the effect of insertions, deletions and duplications.

(A) The type and rate of sequencing errors varies highly among different sequencing platforms (*71*). In the case of the most popular short-read platform, Illumina, sequencing errors typically lead to substitutions at the rate around 0.1% per base pair (*71*). Such a substitution could occasionally lead to a HM, if it occurs at a SNV site and the erroneously called base pair by chance matches the other SNV allele.

(B) Some types of errors in the sequence of a diploid genome may lead to HM. The possible cases are:

(B1) an error at a site without a SNV (e.g. the true sequence is A in both alleles, which we wrote as A-A, but in the genome database it is T-T) - in this case, reads coming from either homolog would contain a mismatch with the database. Such error does not lead to HM.

(B2) a missing SNV (e.g. true A-T, but in the database it is A-A). Such an error leads to mapping imbalance, *i.e.* reads from the homolog with the correct sequence would get mapped, while the reads from a homolog with a sequence error will be dropped. A missing SNV does not lead to HM.

(B3) a false positive SNV (e.g. true A-A, database A-T). In this case, reads from both homologs will be assigned to the same allele (the one matching the true sequence). A false positive SNV error leads to both HM and mapping imbalance.

(B4) an erroneous SNV (e.g. true A-T, database A-C). In this case, reads from the homolog containing an error will not be mapped, which will produce mapping imbalance. An erroneous SNV does not lead to HM.

(B5) missing heterozygosity of an SNV allele. Samples derived from populations of organisms can contain heterozygous SNVs, *i.e.* different organisms within the population may contain different base pairs at the same location on the same homolog. The issue, however, is that genomic databases can store only one allele per homolog (below, referred to as a “known” allele), while the other alleles become ignored (referred to as “unknown” alleles) (note that this issue can be solved by using graph genomes, which can store multiple overlapping variants per position (*72*)). The effects of heterozygosity on mapping depends on the nature of “known” and “unknown” alleles on the two homologs:

(B5.1) non-overlapping “known” and “unknown” alleles, when all alleles of one homolog are different from the allele on the other homolog (e.g., true A/C-T, database A-T). This scenario leads to a partial mapping imbalance, because the reads containing the “unknown” allele do not get mapped. This does not lead to HM.

(B5.2) overlapping “known” and “unknown” alleles, when the “unknown” allele on one homolog is the same as the “known” allele on the other homolog (e.g. true A/T-T, database A-T). This scenario leads to HM, because all reads containing the “unknown” allele will be interpreted as containing the “known” allele from the other homolog. Also, this scenario leads to a mapping imbalance.

#### *Trans*-homolog contacts can be contaminated by *cis* contacts with a misassigned homolog

HM is particularly problematic for studying the *trans*-homolog contact patterns. If we would assign a wrong homolog to one side of a *cis* Hi-C pair, we would misinterpret this pair as a *trans*-homolog contact.

Given that *trans*-homolog contacts in our sample were much less frequent than *cis* ones, even a small probability of HM could lead to a significant contamination of the *trans*-homolog contact map. Note, that a cross-contamination between the *cis* maps of the two homologs was much less likely, since it required simultaneous HM on both sides of a Hi-C pair.

#### Estimating the level of homolog misassignment using Hi-C byproducts

The observed *trans*-homolog read pairs thus was a mixture of “true” *trans*-homolog contacts and *cis* pairs with a misassigned homolog on one side. For P(s) curves, this could be expressed as:

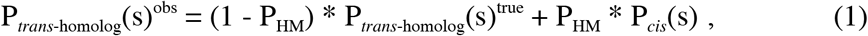

where P_*trans*-homolog_(s)^obs^ was the P(s) curve for the observed *trans*-homolog contacts, P_*trans*-homolog_(s)^true^ was that for the *true trans*-homolog contacts (*i.e.* Hi-C *trans*-homolog pairs that represented a true *trans*-homolog ligation) and P_*cis*_(s) was P(s) for *cis* contacts, and P_HM_ was the probability of HM. Here, the probability of HM was defined per paired read (*i.e.* the probability per side is a P_HM_/2); we assumed that each Hi-C pair had the same chance to get an erroneously assigned homolog, regardless of its genomic separation or orientation, *i.e.* that HM probability was the same for all Hi-C pairs. We also accounted for HM in *trans*-homolog read pairs, which were falsely interpreted as *cis* ones.

Estimating the probability of HM in a mapped Hi-C library was a challenging task. A priori, for any given *trans*-homolog read pair, we cannot tell if it was produced by a true *trans*-homolog ligation, or it resulted from HM of a *cis* pair. However, we could reduce the relative amount of homolog-misassigned pairs by increasing the stringency of homolog assignment. For that purpose, we selected pairs where both sides overlapped at least two or three SNVs and where read alignments had no mismatches with respect to the reference genome. We reasoned that, for such pairs, HM was less likely, since it required simultaneous errors at multiple SNVs.

For pairs with 2+ SNVs on each side, the shape of P_*trans*-homolog_(s) changed at separations <1 kb: the ~10-fold enrichment of *trans*-homolog inward pairs at s<1 kb that was present in unfiltered data (fig. S4A), was greatly reduced after increasing the stringency of homolog assignment (Fig. 2C and fig. S4B). We tried increasing the stringency of homolog assignment further by requiring 3+ SNVs on each side, but this did not change the shape of P_*trans*-homolog_(s) (fig. S4C). This observation suggested that the homolog-misassigned *cis* pairs did not significantly contribute to P_*trans*-homolog_(s) for pairs with 2+ SNVs. Conversely, decreasing the accuracy of homolog assignment by remapping our data to less accurate DGRP SNV annotations (17), increased the abundance of observed short-distance inward *trans*-homolog pairs (fig. S4D). We concluded that the enrichment of short-distance inward *trans*-homolog pairs was due to homolog-misassigned inward *cis* pairs.

Inward *trans*-homolog pairs at s<1 kb were particularly sensitive to contamination by homolog-misassigned *cis* pairs, because at these separations and directionalities *cis* pairs were the most abundant (>10^4^ more abundant that *trans*-homolog pairs of other directionalities) due to “dangling ends”. Thus, even rare HM at a probability as low as ~10^-4^ would generate enough false *trans*-homolog pairs to match the true signal, and thus change the shape of inward P_*trans*-homolog_(s) at s<1 kb. Thus, the ratio of P_*trans*-homolog_(s)/P_*cis*_(s) for inward pairs at short separations could be used as an upper estimate for the probability of HM:

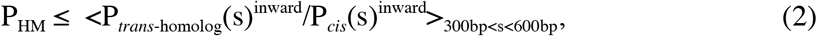

where <…>_300bp<s<600bp_ was the geometric mean around the range of separations between 300 bp and 600 bp, corresponding to the “dangling end” peak of P_*cis*_(s).

Using this approach, we estimated the HM probability in our data at ~0.17% (Fig. 2D). Importantly, this was an *upper* estimate, since not all of the *trans*-homolog inward pairs at s<1 kb were homolog-misassigned *cis* pairs. Finally, given this probability, misassigned *cis* contacts (the second term in Eq. (1)) made up only 5% of detected *trans*-homolog contacts at s~1 kb separations (Fig. 2E) and even less at larger separations.

#### Analysis of published haplotype-resolved mouse Hi-C data

We tested our approach to haplotype-resolved Hi-C using data from two published haplotype-resolved Hi-C studies on mice (*26*).

For mapping, we reconstructed two sets of diploid genomes (Black6 x DBA/2J and Black6 x PWK/PhJ) using SNVs from the Mouse Genome Project (bioRxiv (*73*)). The genome of Black6 parental mouse line did not require reconstruction, since it served as the basis for the reference mm10/GRCm38 genome. We reconstructed consensus genomes with ‘bcftools consensus’, using only homozygous SNVs that passed the quality control. We then mapped the publicly available sequenced Hi-C libraries to the diploid genomes (Black6 x DBA/2J for (*26*) and Black6 x PWK/PhJ for (*25*)) and extracted the positions of Hi-C contacts using the same approach as for our *Drosophila* data. For contact maps, we only used Hi-C pairs overlapping 2+ SNVs on each side. We did not perform IC on the produced contacts maps due to their sparsity. Non-ICed raw contact maps have to be interpreted with a degree of caution, since they are affected by the variation of sequencing visibility of loci due to an uneven distribution of SNVs, GC content, amount of open chromatin, etc.

In a naive interpretation, a decay of *trans*-homolog contact frequency P_*trans*-homolog_(s) with distance s is indicative of homolog pairing, since it means that pairs of loci in homologous positions make more contacts than loci in non-homologous positions. However, such decay can also be observed in samples without pairing, but due to severe contamination of *trans*-homolog contacts by homolog-misassigned *cis* pairs. To interpret the curves of *trans*-homolog contact frequency P_*trans*-homolog_(s), we compared them to a negative control, no-pairing curves P_*trans*-homolog_(s)^no-pairing^, which described the expected amount of observed *trans*-homolog contacts in samples without pairing, but in presence of HM. P_*trans*-homolog_(s)^no-pairing^ could be derived from Eq.(1) assuming P_*trans*-homolog_(s)^true^=const:

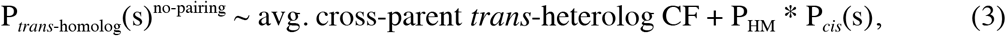

here, we could estimate P_HM_ using Eq. (2).

Thus, in order to claim that a sample had homolog pairing, we had to observe significantly more contacts than that predicted by P_*trans*-homolog_(s)^no-pairing^, at least in some range of separations.

#### Analysis of published Hi-C data from *Drosophila* embryos

To compare our data with other studies on the structure of chromosomes in *Drosophila* embryos, we re-analysed the publicly available Hi-C dataset from (*38*) using the same methods as we used for our own data.

#### Pairing score

To characterize the degree of pairing between homologous loci across the whole genome, we introduced a genome-wide statistics track called *pairing score (PS)*. The PS of a genomic bin is log_2_ of average *trans*-homolog IC contact frequency between all pairs of bins within a window of +-W bins. For each genomic bin i, its pairing score with window size W was defined as:

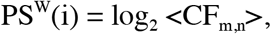

averaged over bins *m* and *n* between i-W-th and i+W-th genomic bins on different homologs of the same chromosome.

Importantly, under this definition, the PS quantified only contacts between homologous loci and their close neighbors and did not quantify pairing between non-homologous loci on homologous chromosomes.

The choice of the window size W was guided by the balance between specificity and sensitivity. Using a bigger window increased sensitivity, accumulating contacts across more loci pairs, while smaller windows increased specificity, allowing to see smaller-scale variation of homolog pairing; empirically, we found that, for our contact maps binned at 4 kb resolution, using a 7×7 bin window (W=3) provided the optimal balance between specificity and sensitivity.

A close examination of the PS track revealed two important features: (i) the *Drosophila* genome was divided into regions that demonstrate consistently high, relatively similar, values of PS, followed by extended regions where PS dipped into lower values, (ii) switching between high- and low-PS regions seemed to occur around insulating boundaries.

#### Insulation scores

The tracks of contact insulation score were calculated using the package cooltools insulation (https://github.com/mirnylab/cooltools). The method used in the package was based on the algorithm described in (*74*), and modified in (*75*).

For every bin of a contact map binned at 4 kb resolution, we calculated the insulation score as the total number of normalized and filtered contacts formed across that bin by pairs of bins located on the either side, up to 5 bins (20 kb) away for Hi-C mapped to reference dm3 maps, and up to 10 bins (40 kb) away for haplotype-resolved Hi-C maps. We then normalized the score by its genome-wide median. To find insulating boundaries, we detected all local minima and maxima in the log_2_-transformed and then characterized them by their prominence (Billauer E. peakdet: Peak detection using MATLAB, http://billauer.co.il/peakdet.html). The detected minima in the insulation score corresponded to a local depletion of contacts across the genomic bin, were then called as insulating boundaries. We found empirically that the distribution of log-prominence of boundaries had a bimodal shape, and we selected all boundaries in the high-prominence mode above a prominence cutoff of 0.1 for Hi-C mapped to the reference dm3 map, and a cutoff of 0.3 for haplotype-resolved Hi-C maps. We called the insulating boundaries in *trans*-homolog contact maps using the same technique, requiring a minimal prominence of 0.3. Finally, we removed boundaries that were adjacent to the genomic bins that were masked out during IC.

To estimate the similarity of insulating boundaries detected in the *cis* and *trans*-homolog contact maps, we calculated the number of overlapping boundaries. We allowed for a mismatch up to 4 genomic 4 kb bins (16 kb total) between overlapping boundaries to account for the drift caused by the stochasticity of contact maps. We reported average percentage of overlapping boundaries. For instance, the average fraction (73.9%) that overlapping boundaries between *cis* maternal and *cis* paternal boundaries occupied within *cis* maternal only (74.2%), and *cis* paternal only boundaries (73.7%). Similarly, for replicates, we calculated the average percentage of overlapping boundaries for *cis* maternal replicates (75.4%), and also for *cis* paternal replicates (71.1%), and then provided a mean of the two values (73.2%). In reference dm3 Hi-C contact maps, the percentage of overlapping boundaries between the two replicates reached 92.3%, suggesting that the lower reproducibility of haplotype-resolved boundaries is due to technical and not biological variation between the two replicates.

We estimated the significance of an overlap between two given boundary sets by comparing it to the overlap expected by chance alone, given the sizes of the two sets. We estimated the latter by calculating an overlap between two similarly-sized random subsets of genomic bins visible in our Hi-C maps. To estimate how significantly the observed overlap deviates from the random expectation, we repeated randomization 10 times and compared the random overlaps to the observed overlap using a t-test.

#### ChIP-seq

We mapped the publicly available raw ChIP-seq data following the same procedure as used by the ENCODE consortium (*76*), (https://github.com/ENCODE-DCC/chip-seq-pipeline).

To generate the tracks of Zld binding, we used ChIP datasets GSM1596215 and GSM1596219 and input datasets GSM1596216 and GSM1596220 from study GSE65441 (*50*). For Dl, we used ChIP datasets GSM1596223 and GSM1596227 and input datasets GSM1596224 and GSM1596228 from the same study GSE65441 (*50*). For GAF, we used ChIP datasets GSM614652 and input datasets GSM614653 and GSM614654 from study GSE23537 (*53*). Finally, for Bcd, we used ChIP datasets GSM1332670 and GSM1332671 and input datasets GSM1332672 and GSM1332673 from GSE55256 (*48*).

#### Correlation analyses between the pairing score and ChIP signal

The correlation analyses were performed as previously described (*77*). The genome was divided into 10 kb bins. In each bin, the fraction of sequence occupied by the PS or the respective ChIP signal was determined. Then, for each bin genome-wide correlations were calculated between the PS and a ChIP signal of interest. The strength of correlation was reported using pairwise Spearman correlation coefficients and corresponding P-values. The Spearman correlation coefficients were also displayed using a heatmap. Control regions consisted of randomized 200 kb chunks of the pairing score.

#### Correlation between the pairing score and Zld binding

To visualize the relationship between PS and Zld binding, we plotted the distribution of PS for genomic bins from each of the four quartiles of the genome-wide distribution of Zld ChIP-seq signal.

We then tested if the effects of Zld binding on chromatin conformation were correlated with the pairing status of loci. We re-analyzed the published Hi-C datasets on Zld depletion (*38*) with the same computational methods as above and obtained normalized Hi-C maps at 4 kb resolution for WT and Zld-depleted embryos at nuclear cycle 14. Then, we calculated the change of the 20 kb-window insulation score upon Zld depletion, Δins^Zld^, and compared it to PS. For bins from each quartile of Zld ChIP-seq binding, we fitted the resulting 2D-distribution of Δins^Zld^ *vs* PS with a bivariate Gaussian. We then plotted the resulting Gaussians as ellipses corresponding to +-2 standard deviations from the mean along each of the independent axes of the distribution (*i.e.* Mahalanobis distance of 2).

#### Other

We performed all custom data analyses in Jupyter Notebooks (*78*), using matplotlib (*79*), numpy (*80*), and pandas (*81*) packages. We automated data analyses in command line interface using GNU Parallel (*82*).

---

**Fig. S1.**
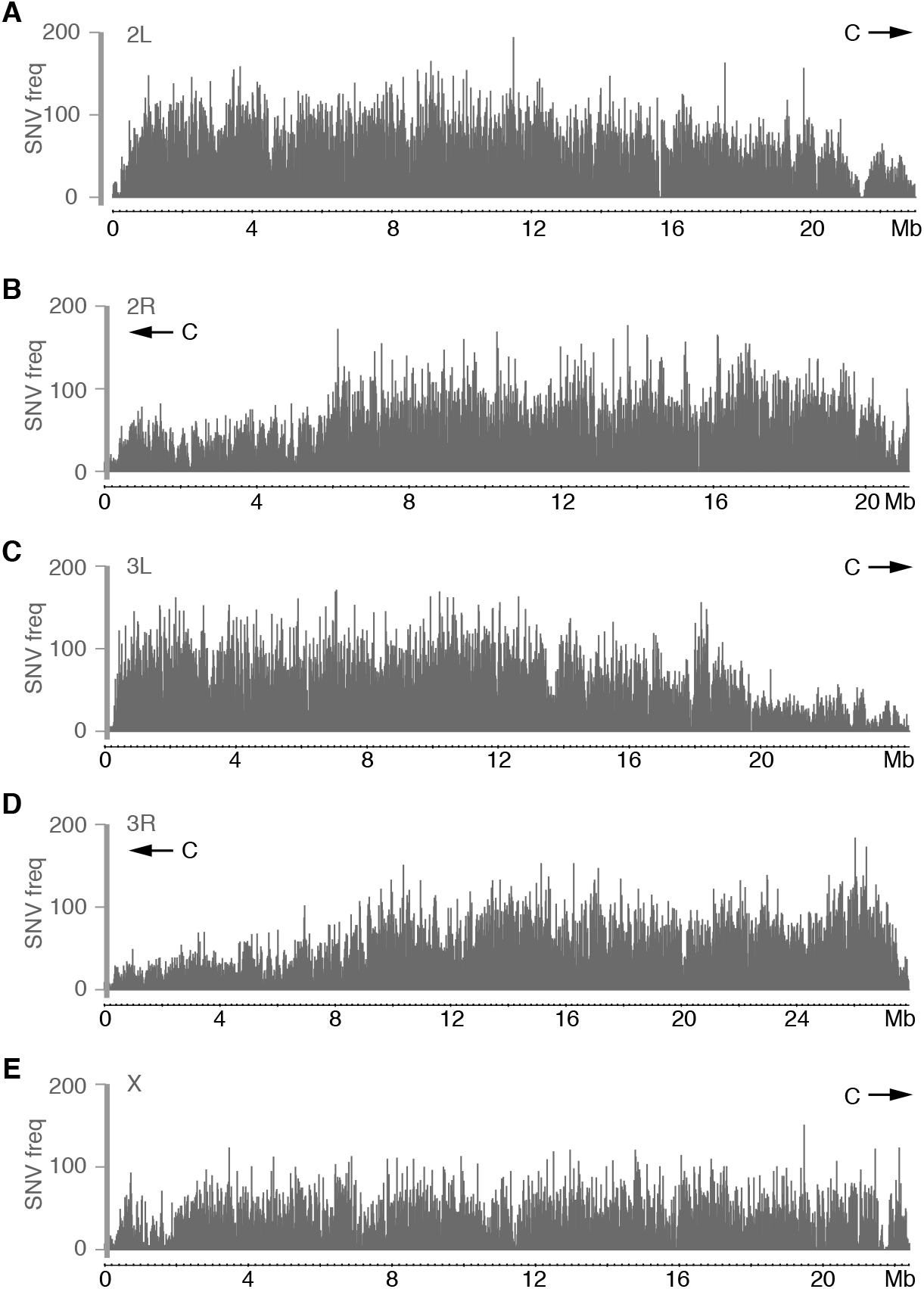
Distribution of SNVs in the F1 hybrid *Drosophila*. (**A** to **E**) The distribution of SNVs along 2L (**A**), 2R (**B**), 3L (**C**), 3R (**D**), and X (**E**) at 4 kb resolution. Note that the SNV frequency decreases in pericentromeric regions. Arrow points towards direction where centromere (C) should be located.

**Fig. S2.**
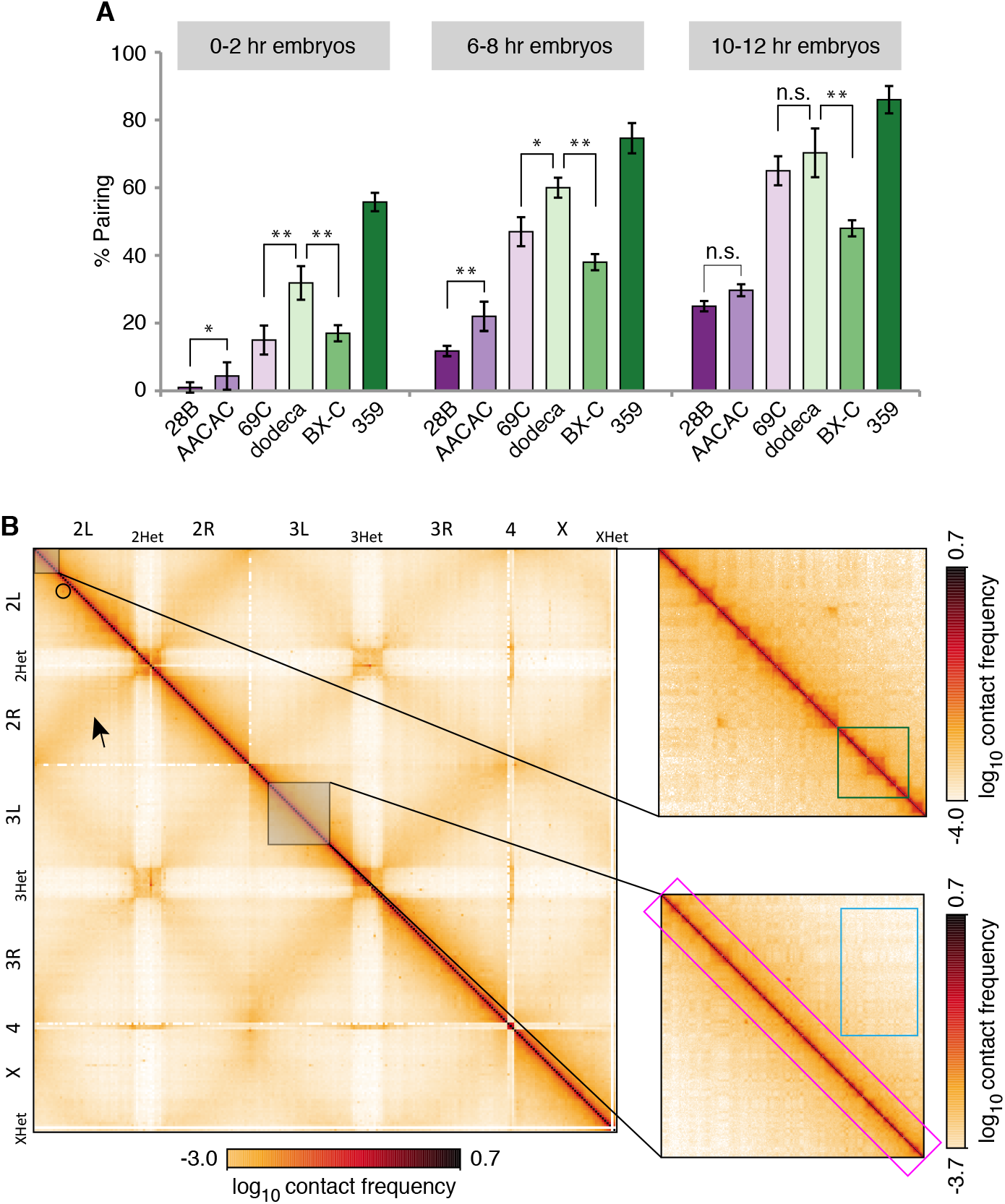
Assessment of homolog pairing by FISH, and of Hi-C data quality in the F1 hybrid embryos. (**A**) Levels of homolog pairing increase steadily as development progresses in the F1 hybrid embryos. Percentage of nuclei showing paired loci at the FISH targets from Fig. 2D in 0-2 hr, 6-8 hr, and 10-12 hr embryos (error bars, standard deviation of at least 3 replicates; n ≥100 nuclei/replicate). Nuclei are considered paired nuclei when FISH signals are ≤0.8 μm (center-to-center distance) apart. Pairing levels at heterochromatic targets are significantly higher than at euchromatic that lie on the same chromosome during 0-2 hr, 2-4 hr (Fig. 2E), and 6-8 hr. However, later during development, at 10-12 hr, levels become more similar (*P<0.05, **P<0.0001, n.s. not significant; Fisher’s two-tailed exact). (**B**) Genome-wide Hi-C map of the 2-4 hr F1 hybrid embryos mapped to the reference dm3 genome. Greyed out boxes, zoomed-in views on 2L and 3L. Features such as central *cis* diagonal (pink box), domains (green box), plaid-patterned compartments (blue box), interaction peaks (circle), and contacts consistent with Rabl (arrow) are indicated. The maps were visualized using the HiGlass browser (*83*).

**Fig. S3.**
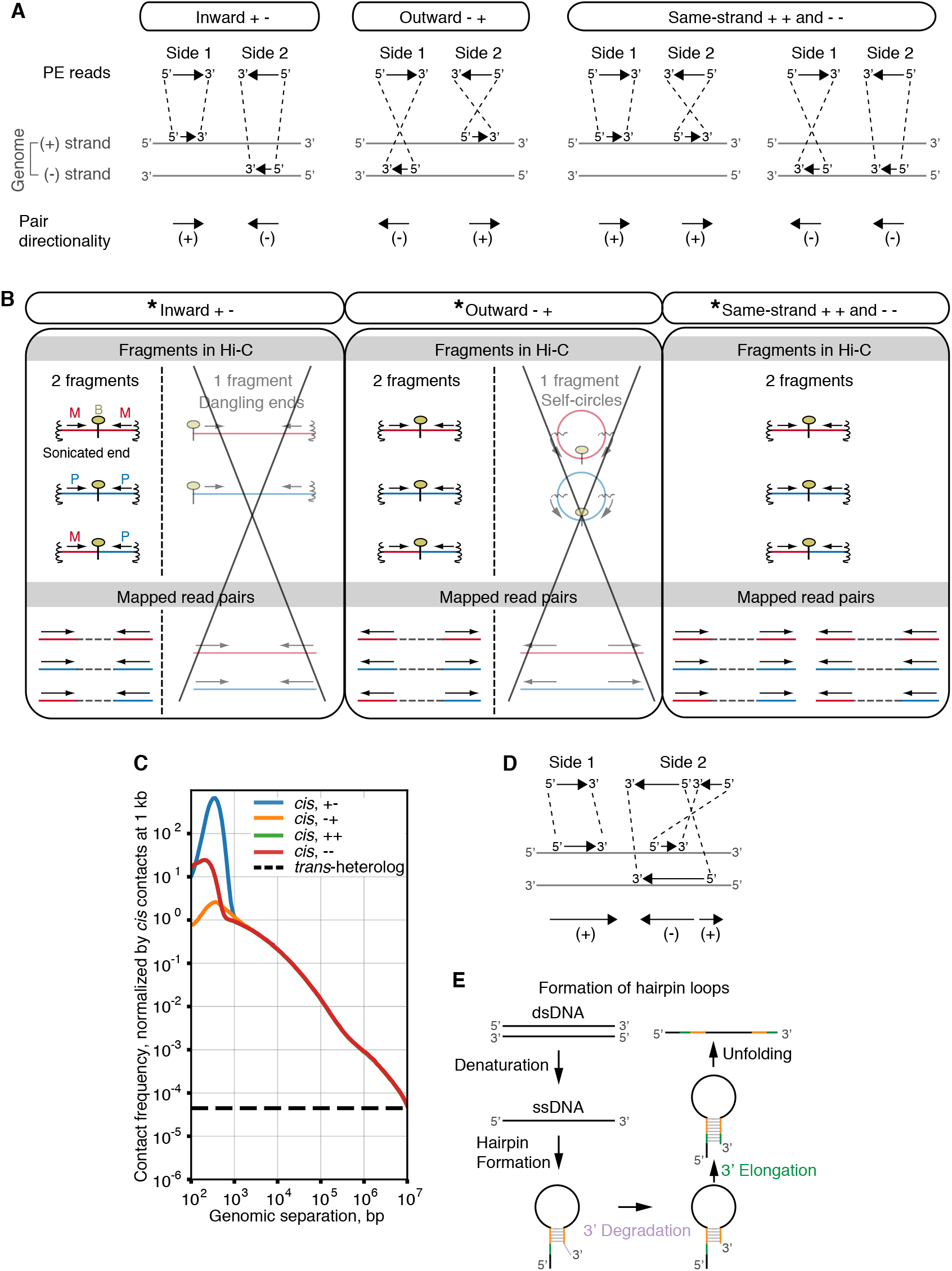
The classification of Hi-C byproducts. (**A**) Mapped read pairs can be classified based on relative locations and directions of both sides of a read pair into three types: inward, outward, and same-strand. (**B**) As adapted from (*44*), Hi-C byproducts manifest themselves as Hi-C pairs of a specific directionality. These byproducts appear as inward or outward pairs, and should not appear in *trans* (crossed out) (asterisk, see supplementary materials and methods for other sources of contamination). Red, maternal (M); blue, paternal (P); yellow ellipse, biotin; wiggly vertical line, sonicated ends. (**C**) Contact frequency plotted against genome separation using ≤ 1 SNV per read, and split by read pair directionality into inward, outward, and same-strand. (**D**) A common mapping pattern of read pairs observed in our data, with one read fully aligned to one genomic locus, and the other side split into two alignments of opposite directionality. (**E**) Formation of hairpin loops, as suggested to be the source of contamination among same-strand reads (*68, 69*). Upon denaturation, palindromic sequence of ssDNA (orange) may form hairpin structure. Unannealed 3’ ends (purple) may be degraded, and refilled with sequence based on 5’end (green) during generation of the Hi-C library.

**Fig. S4.**
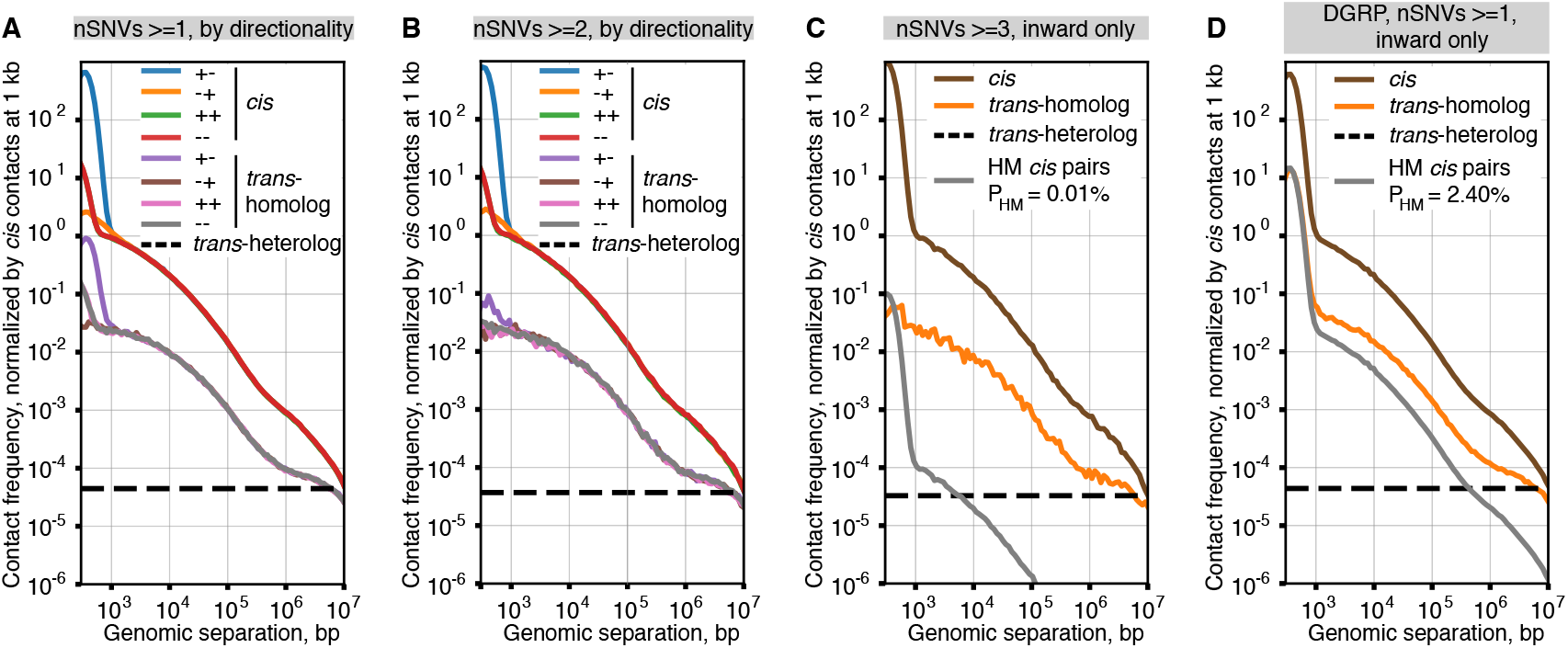
The rate of homolog misassignment depends on the number and quality of SNVs used for alignment. (**A** and **B**) Contact frequency plotted against genome separation, and split by read pair directionality to inward, outward, and same-strand for pairs containing at least (**A**) 1 SNV or (**B**) 2 SNVs per read. Requiring ≤2 SNVs per read removed a characteristic enrichment of inward *trans*-homolog contact frequency at s<1 kb, which suggested that this enrichment in unfiltered data was caused by homolog misassignment (HM) of *cis* pairs. (**C**) Contact frequency for inward read pairs with at least 3 SNVs, with no sequence mismatches allowed (P_HM_=0.01%). (**D**) Contact frequency for inward read pairs mapped using the DGRP SNV annotations (*39*) requiring at least 1 SNV per read (P_HM_=2.4%). (**A** to **D**) Contact frequencies for chromosomes 2 and 3 normalized by *cis* contact frequency at 1 kb. Dashed black line, average *trans-*heterolog contact frequency; P_Hm_, probability of homolog misassignment.

**Fig. S5.**
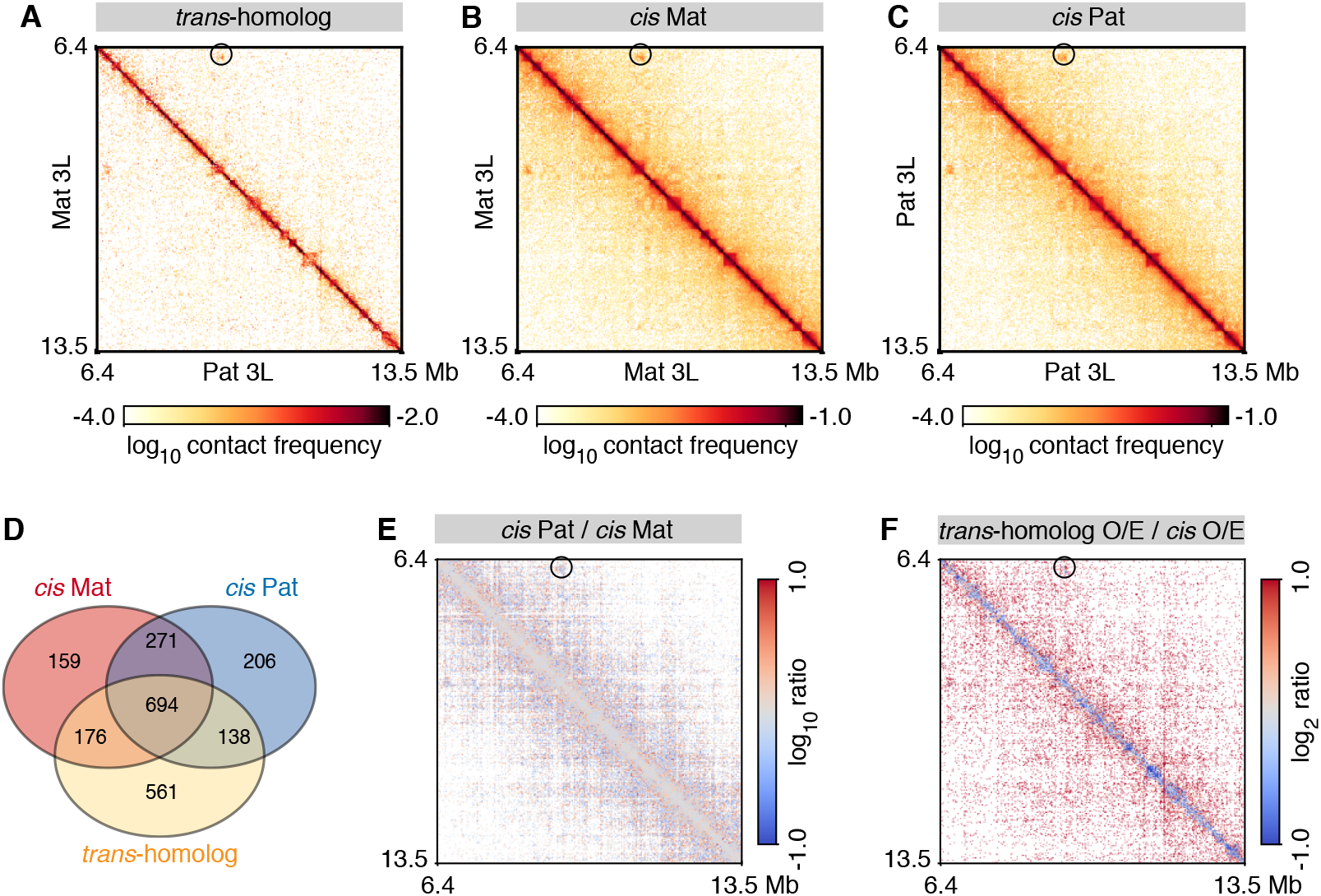
Haplotype-resolved Hi-C maps suggest highly structured *trans*-homolog domains, boundaries, and interaction peaks that resemble analogous *cis* features. (**A**) *Trans*-homolog, (**B**) *cis* maternal, and (**C**) *cis* paternal maps of matching ~7 Mb regions on 3L. (**D**) Venn diagram shows the extent of overlap among *trans*-homolog boundaries, maternal and paternal *cis*-boundaries. (**E**). The ratio of *cis* Pat/*cis* Mat Hi-C maps indicates the *cis* contact patterns of two homologs are highly concordant. (**F**) The ratio of *trans*-homolog/average *cis* maps suggests that pairing resembles *cis* contacts, albeit with lower interactions in some regions (dark blue). (**A**, **B**, **C**, **E**, and **F**) Interaction peaks are encircled. The maps displayed (3L:6.4-13.5Mb) represent a zoomed-out from the maps (3L:9.85-11.6Mb) in Fig. 3I to M.

**Fig. S6.**
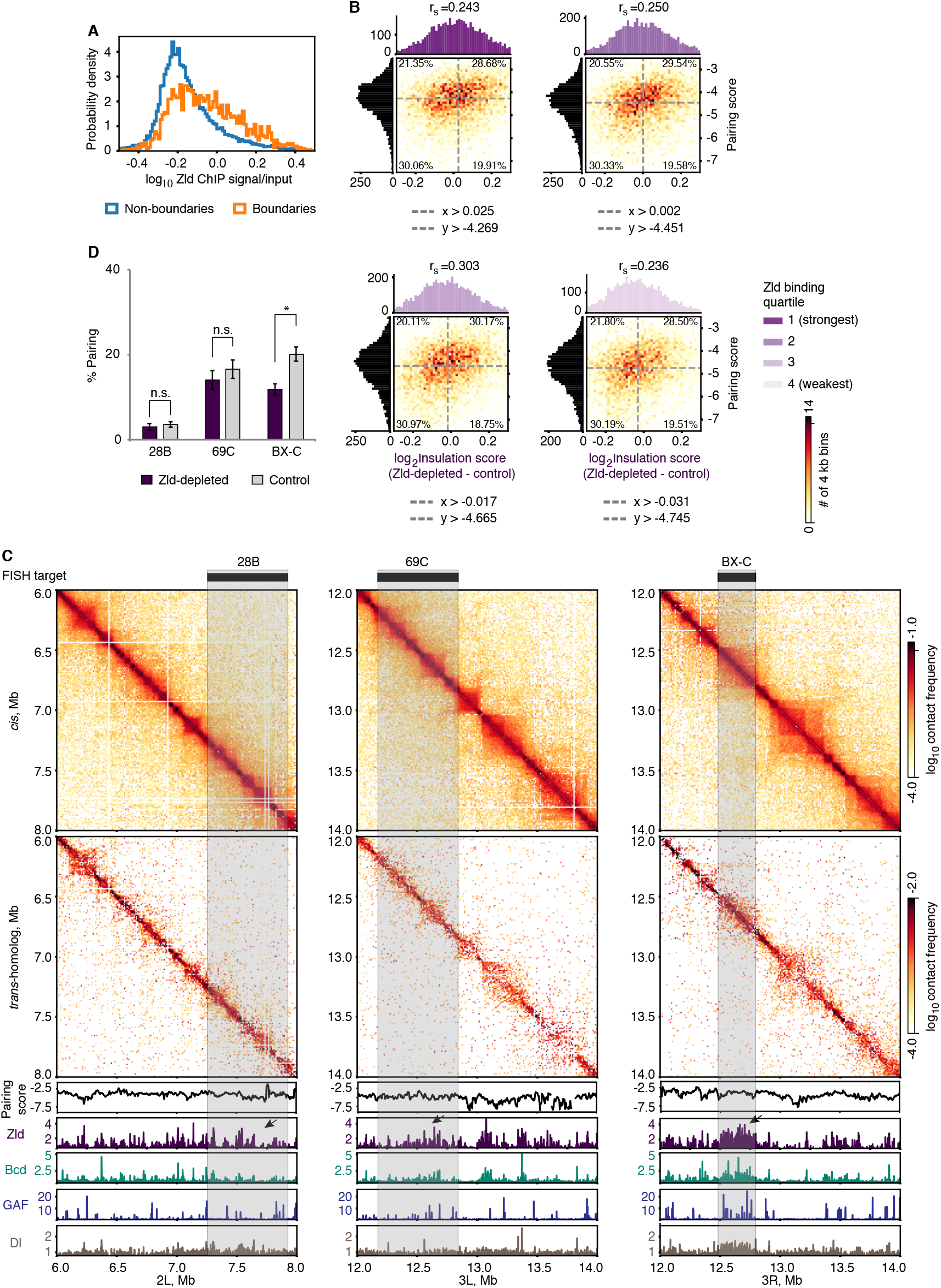
Zld-dependent boundary formation is associated with homolog pairing. (**A**) Distribution of Zld signal suggests that domain boundaries are more likely to have strong Zld binding than non-boundaries (P<10^-10^, Mood’s median test). (**B**) Distribution of the difference in insulation scores between Zld-depleted and control nc14 embryos relative to the pairing score, stratified by the Zld binding quartiles (also depicted as fitted ellipses in Fig. 4D). Dashed lines, medians; r_s_, Spearman’s correlation coefficient. (**C**) *Cis* (upper panels) and *trans*-homolog (second panels from the top) contact maps at 2L:6-8 Mb, 3L:12-14 Mb, and 3R:12-14 Mb. Lower panels, pairing score (PS) calculated using a 28 kb window at 4 kb resolution (black), the ChIP-seq profiles of Zld (dark purple) (*50*), Bcd (green) (*48*), GAF (blue) (*53*), and Dl (brown) (*50*). Grey boxes, locations of FISH targets; arrows, Zld signal at FISH targets. (**D**) Percentage of nuclei showing paired loci for Zld-depleted and control embryos (cycle 14; error bars, standard deviation of at least 4 replicates; n ≥100 nuclei/replicate; *, P = 2.56×10^-3^, n.s. not significant, Fisher’s two-tailed exact).

**Fig. S7.**
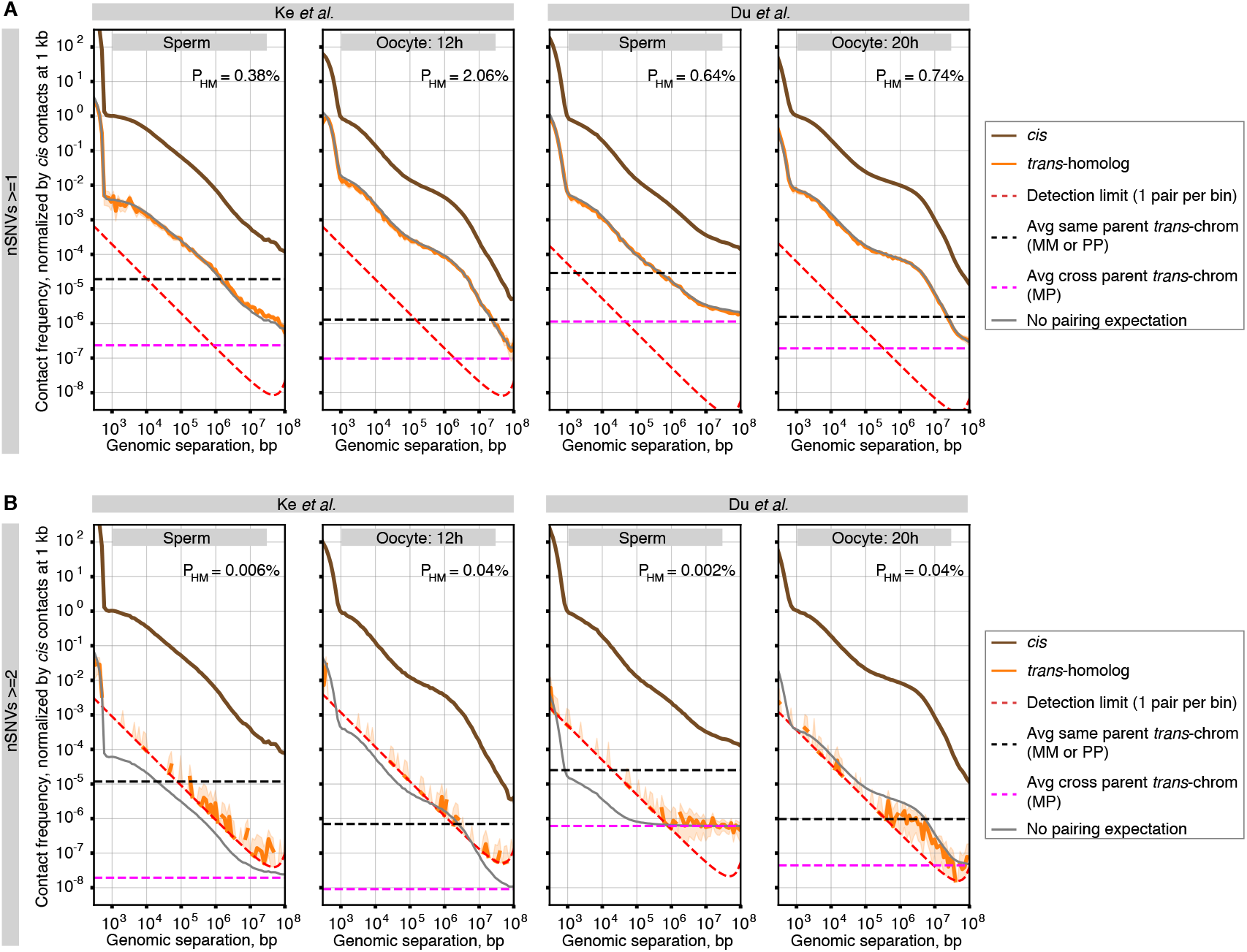
A stringent filtering of haplotype-resolved Hi-C datasets from mouse gametes removes *trans*-homolog pairs. (**A** and **B**) Contact frequency plotted against genome separation in Ke *et al.* (*26*) and Du *et al.* (*25*) gamete Hi-C datasets, requiring at least (**A**) 1 or (**B**) 2 SNVs per read. Shaded area, 95% confidence intervals calculated via Poisson resampling. Dotted lines, average *trans* contact frequency as a function of distance between the same parent of origin chromosomes (black; MM or PP), or between maternal and paternal chromosomes (pink; MP). Grey line, expected *trans*-homolog contact frequency as a function of distance without pairing, but in presence of homolog misassignment (HM). No sequence mismatches allowed for all read pairs. Maternal (M), and paternal (P) chromosomes. P_Hm_, probability of homolog misassignment.

**Fig. S8.**
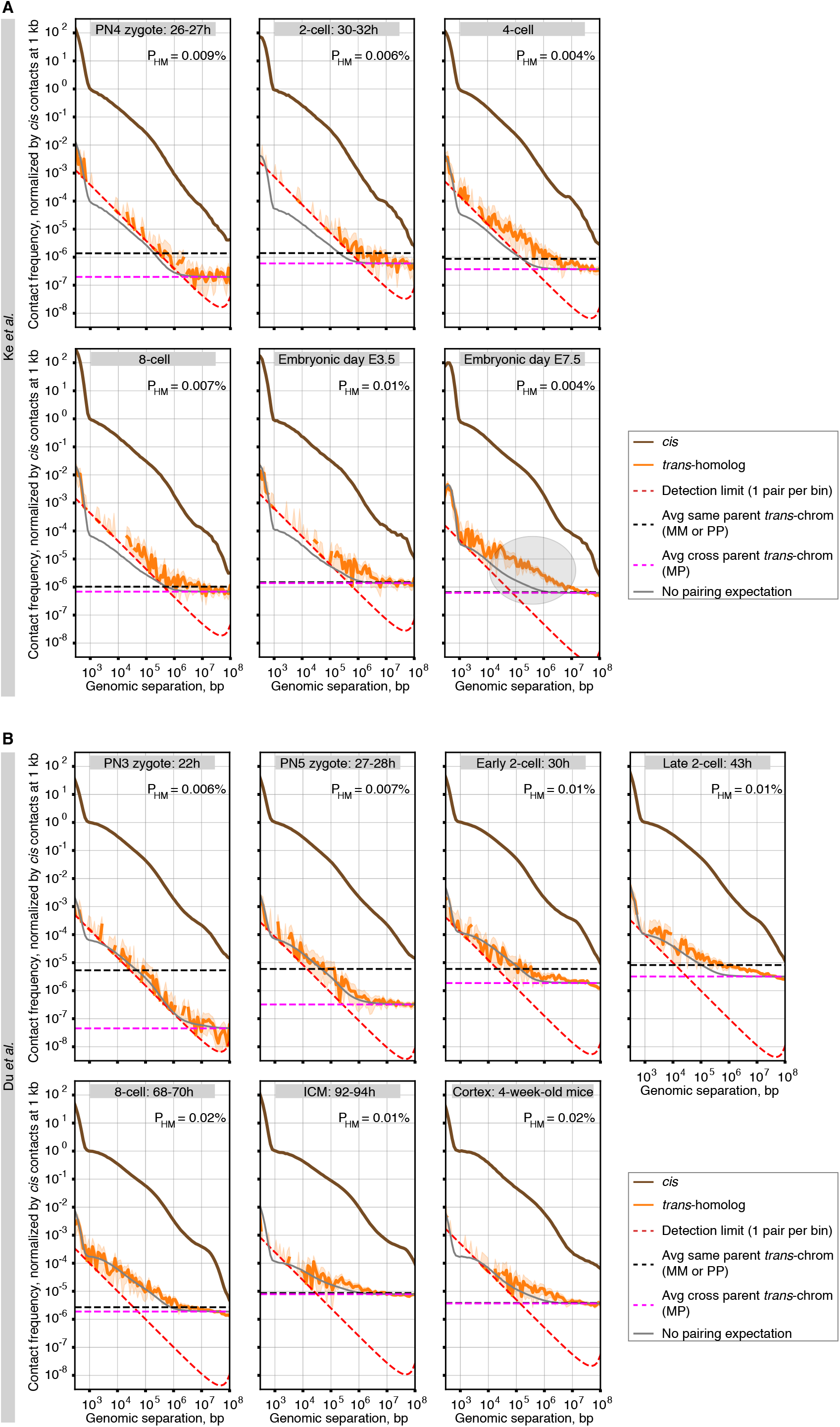
Haplotype-resolved Hi-C datasets representing hybrid mouse embryos show a minor population of *trans*-homolog read pairs. (**A** and **B**) Contact frequency plotted against genome separation in (**A**) Ke *et al.* (*26*) or (**B**) Du *et al.* (*25*) datasets, requiring at least 2 SNVs per read. Shaded area, 95% confidence intervals calculated via Poisson resampling. Dotted lines, average *trans* contact frequency as a function of distance between the same parent of origin chromosomes (black; MM or PP), or between maternal and paternal chromosomes (pink; MP). Grey line, expected *trans*-homolog contact frequency as a function of distance without pairing, but in presence of homolog misassignment (HM; no pairing expectation). No mismatches allowed for all read pairs. Maternal (M), and paternal (P) chromosomes. The grey ellipse (E7.5) highlights the area where *trans*-homolog contacts exceed the no pairing expectation. P_Hm_, probability of homolog misassignment.

**Fig. S9.**
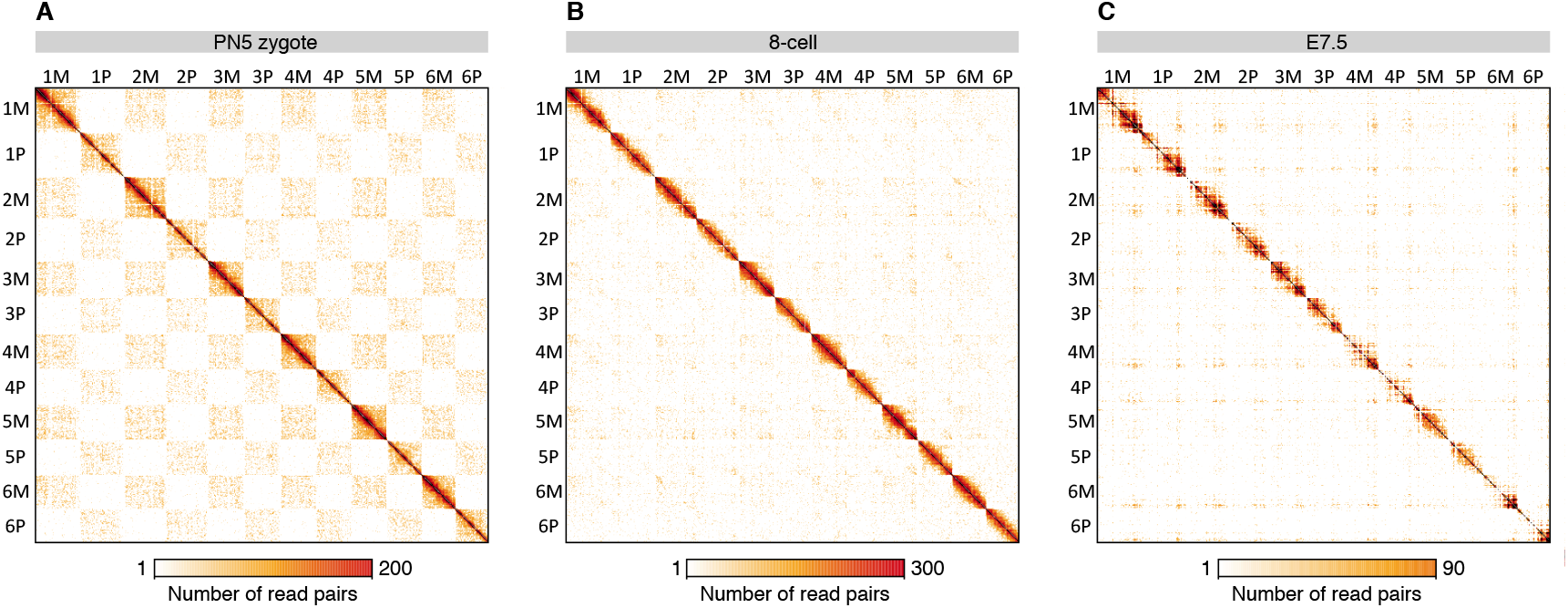
Segregation of parental genomes reduces during developmental progression. (**A** to **C**) Haplotype-resolved Hi-C maps with 2 SNVs per read are displayed using the HiGlass browser (*83*) for (**A**) PN5 zygote and (**B**) 8-cell embryos from Du *et al.* (*25*), and (**C**) E7.5 embryos from Ke *et al.* (*26*). Maternal (M); paternal (P); M, C57BL/6N; P, PWK/PhJ for Du *et al.* datasets, while M, C57BL/6J; P, DBA/2J mice strain for Ke *et al.* dataset.

**Table S1.** Variant annotations for maternal (DGRP-057) and paternal (DGRP-439) lines. Numbers of indels and SNVs are shown for each chromosome in both homozygous (Hom) and heterozygous (Het) scenario. Maternal (Mat); paternal (Pat).

| Chrom Name | Chrom Size | Indels |  |  | SNVs |  |  |  |
| --- | --- | --- | --- | --- | --- | --- | --- | --- |
|  |  | Hom | HetMat | HetPat | Hom | HetMat | HetPat | Het per kb |
| **2L** | 23,011,544 | 9,415 | 5,820 | 5,910 | 68,091 | 66,894 | 70,989 | 5.992 |
| **2R** | 21,146,708 | 8,828 | 5,144 | 5,396 | 60,606 | 55,659 | 58,689 | 5.407 |
| **3L** | 24,543,557 | 10,374 | 6,068 | 5,599 | 73,011 | 67,299 | 64,261 | 5.360 |
| **3R** | 27,905,053 | 9,676 | 6,494 | 6,402 | 65,627 | 66,777 | 68,709 | 4.855 |
| **4** | 1,351,857 | 136 | 1 | 4 | 929 | 2 | 15 | 0.013 |
| **X** | 22,422,827 | 5,193 | 8,405 | 8,386 | 38,063 | 40,674 | 44,055 | 3.779 |

**Table S2.** Statistics summary for the dm3 reference mapping and duplicate removal. The number of read pairs is provided for each biological replicate. Total raw read pairs (‘Total’) include unmapped read pairs with multiple or low score alignments (‘Total_unmapped’), and read pairs where only single side (‘Total_ss_mapped’) or both sides are mapped (‘Total_mapped’) (*44*). Total mapped read pairs contain non-duplicates and discarded PCR duplicates. Non-duplicates are subdivided into *cis* and *trans* read pairs. Portion of non-duplicates with respect to total raw read pairs is indicated as mappability.

|  | Replicate 1 | Replicate 2 |
| --- | --- | --- |
| **Total** | 256,988,962 | 256,274,483 |
| **Total_unmapped** | 53,747,071 | 50,506,661 |
| **Total_ss_mapped** | 37,369,722 | 37,037,449 |
| **Total_mapped** | 165,872,169 | 168,730,373 |
| **Duplicates** | 23,641,775 | 21,411,225 |
| **Non-duplicates** | 142,230,394 | 147,319,148 |
| *cis* | 131,681,245 | 134,880,524 |
| *trans* | 10,549,149 | 12,438,624 |
| ***cis/trans* ratio** | 12.5 | 10.8 |
| **Mappability (%)** | 55.3 | 57.5 |

**Table S3.** Statistics summary for the haplotype-resolved mapping and duplicate removal. Haplotype-resolved mapping is performed using SNVs defined in this study or in Mackay *et al.* 2012 (*39*). The number of read pairs is provided for each biological replicate. Total raw read pairs (‘Total’) include unmapped read pairs with multiple or low score alignments comprising also reads without SNVs (‘Total_unmapped’), and read pairs where only single side (‘Total_ss_mapped’) or both sides are mapped (‘Total_mapped’). Total mapped read pairs contain non-duplicates and discarded PCR duplicates. Non-duplicates are subdivided into *cis* and *trans* read pairs. Portion of non-duplicates with respect to total raw read pairs is indicated as mappability. *Given that the number of *trans*-homolog read pairs is 497,766 (Replicate 1) and 528,169 (Replicate 2), 36.1% of all *trans* read pairs are *trans*-homolog.

|  | SNVs from this study |  | SNVs from Mackay *et al.* 2012 |  |
| --- | --- | --- | --- | --- |
|  | Replicate 1 | Replicate 2 | Replicate 1 | Replicate 2 |
| **Total** | 256,988,962 | 256,274,483 | 256,988,962 | 256,274,483 |
| **Total_unmapped** | 183,409,873 | 180,413,502 | 174,706,472 | 171,547,794 |
| **Total_ss_mapped** | 45,654,919 | 47,956,696 | 57,145,342 | 59,638,376 |
| **Total_mapped** | 27,924,171 | 27,904,286 | 25,137,148 | 25,088,313 |
| **Duplicates** | 3,533,675 | 3,245,230 | 2,220,900 | 1,959,943 |
| **Non-duplicates** | 24,390,496 | 24,659,056 | 22,916,248 | 23,128,370 |
| **cis** | 23,076,479 | 23,130,134 | 21,265,473 | 21,243,109 |
| **trans** | *1,314,017 | *1,528,922 | 1,650,775 | 1,885,261 |
| **cis/trans ratio** | 17.6 | 15.1 | 12.9 | 11.3 |
| **Mappability (%)** | 9.5 | 9.6 | 8.9 | 9.0 |

**Table S4.** Statistics summary for the haplotype-resolved mapping and duplicate removal using mammalian data. *See separate excel sheet*. The number of read pairs is provided for each replicate from Ke *et al.* (*26*) and Du *et al.* (*25*) datasets. Total raw read pairs (‘Total’) include unmapped read pairs with multiple or low score alignments comprising also reads without SNVs (‘Total_unmapped’), and read pairs where only single side (‘Total_ss_mapped’) or both sides are mapped (‘Total_mapped’). Total mapped read pairs contain non-duplicates and discarded PCR duplicates. Non-duplicates are subdivided into *cis* and *trans* read pairs.

**Table S5.** Information on Oligopaint probe sets. For each Oligopaint probe set information such as the span in the genome with start and stop coordinates, target size, number of oligos, and probe density are given.

| Probe set | Chr | Start (dm3) | Stop (dm3) | Target size/kb | # Oligos | Probes/kb |
| --- | --- | --- | --- | --- | --- | --- |
| 28B | 2L | 7,256,488 | 7,936,487 | 680 | 10,000 | 14.7 |
| 69C | 3L | 12,170,682 | 12,844,681 | 674 | 10,000 | 14.8 |
| 89D-89E/<br>BX-C | 3R | 12,482,502 | 12,797,965 | 315 | 2,394 | 7.6 |

**Table S6.**
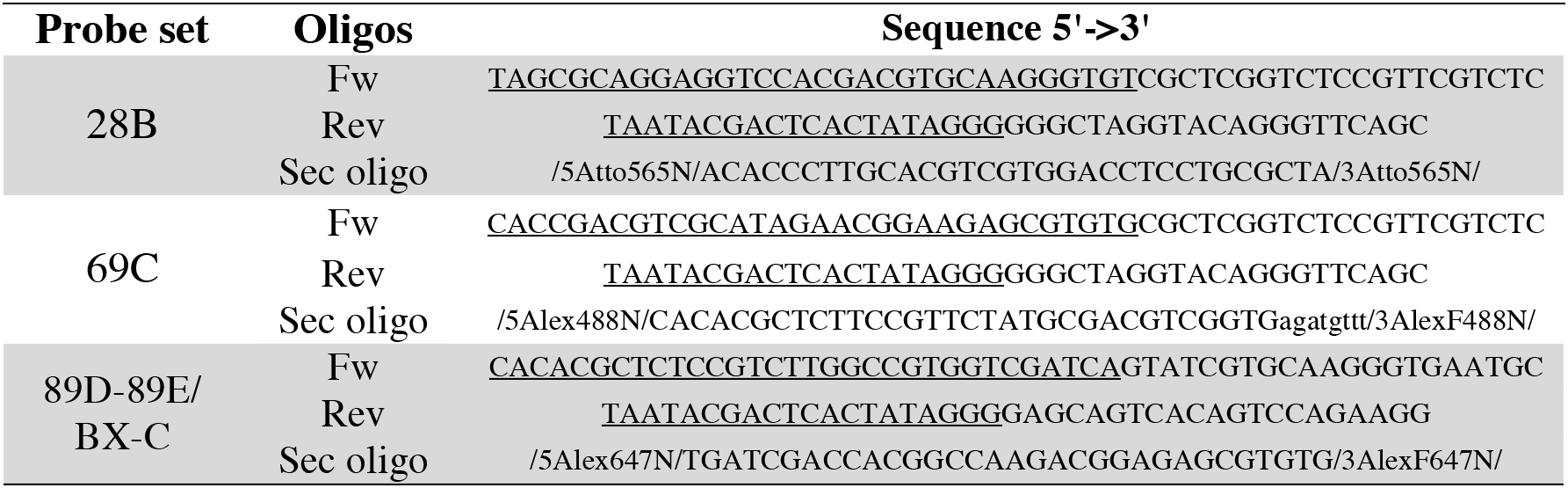
Primers and secondary oligos for Oligopaint probe sets. For each Oligopaint probe set primer and secondary (sec) oligo sequences are given. In forward primer a site for secondary oligo annealing is underlined, while in reverse primer a T7 promoter sequence is underlined. Sequences for secondary oligos (*62*), adapted with a 5’ and 3’ fluorophore of interest, are provided using the modification codes from Integrated DNA Technologies (IDT).

